# Patterns of convergent somatic hypermutations in the adaptive immune response of Mus musculus

**DOI:** 10.1101/2022.10.12.510618

**Authors:** Alexander C. Wenner, Charles A. Mettler, Ellie M. Sharp, Thomas C. Hansen, Isabella B. Vari, Alexander V. Le, Jörg Zimmermann

## Abstract

We analyzed a dataset of 964 clonally unrelated murine antibodies for which structures have been deposited in the PDB. 454 of the 964 antibodies have gapless germline assignments and do not have excessive numbers of computationally identified somatic hypermutations (SHMs). About 5,500 SHMs were identified, of which approximately 3,500 are in the framework. We then searched for correlated convergent SHMs, i.e. groups of SHMs that arose independently in different antibodies but at the same sequence position and with the same germline and mature amino acid identity. A surprisingly large number of groups of correlated convergent SHMs were found. 329 antibodies share at least two, 161 antibodies share at least three, 87 antibodies share at least four, and 53 antibodies share at least five identical SHMs with another antibody in the dataset. We then analyzed whether any of the correlated SHMs are forming structural cluster. Approximately 400 clusters where CFWMs are located within 10 Å of each other were identified. 158 of these clusters are in the framework region. Identification of such structural clusters of correlated convergent SHMs may help identify adaptive mutations that act in an antigen-independent manner.

## Introduction

During a germinal center (GC) reaction, it only takes 5 to 15 somatic hypermutations (SHMs) to turn a naïve germline B-cell receptor (BCR) which has moderate affinity for the cognate antigen and is often polyspecific (including affinity to self-molecules), into a high-affinity binder with exquisite specificity for antigen. [1, 2] Understanding the selection pressures in GC reactions that lead to this outcome and identifying patterns of SHMs has been a longstanding goal of immunological research. [3–5] Many SHMs are in or proximal to the binding site, where they add or optimize specific interactions with antigen. However, a substantial number of SHMs, referred to as framework mutations, are distal from the binding site, and it is often unclear if and how they improve affinity and/or specificity of the BCR. One hypothesis is that framework mutations are stabilizing the variable domain to compensate for destabilizing mutations in the binding site. [3, 6] We and others have hypothesized that framework mutations rigidify the framework and thereby increase antigen-specificity. [7–14] It is also well known that the AID pathway that generates SHMs has strong biases for certain nucleotide motifs, and it has been suggested that some framework mutations are simply the result of neutral genetic drift. [15–21] Regardless of the origin or function of framework mutations, and although they occur at a substantial rate, they are often considered less relevant to the evolution of affinity and specificity during GC reactions. For example, antibody engineering efforts often focus mutational variations on CDR loops, and the framework is left unaltered. [22–28]

The majority of work to date regarding the mutability of antibody framework regions and their roles in structural evolution has been geared towards identifying key positions, residues, and relative orientations of heavy and light chain that modulate binding affinity in specific antibodies. [3, 4, 6, 16, 18, 19, 29] Perhaps the most comprehensive treatment of patterns of SHMs are the so-called gene-specific substitution profiles (GSSPs) developed by Sheng et al.. [29] A GSSP shows the most frequent amino acid substitutions along a given germline V gene. Sheng et al. found that, while common substitutions in a given V gene are largely in line with the codon biases of the replication machinery, the same codon can display different substitution preferences between different V genes, emphasizing the gene-dependence of framework mutations. A possible explanation of this observation is that common framework mutations are in fact independent of ligand insofar as the corresponding V genes are a germline precursor for specific “ligand types” (e.g., peptide, small organic, polysaccharide, etc.). Common mutations could then represent a step towards convergent structural evolution unique to the V gene. This may also explain why V gene repertoires are highly similar in sequence in their framework regions. [30]

Here we use a dataset of 454 antibodies for which structures have been deposited in the Protein Data Bank to identify structural clusters of convergent framework SHMs. We hypothesize that such mutations are more likely to be instances of adaptive, antigen-independent mutations because (i) framework mutations are not in direct contact to antigen, thus they are more likely to act in an antigen-independent manner, (ii) convergent SHMs are mutations that arose independently in different individuals, hence, they are more likely to be adaptive since it is less likely that the same neutral mutations occur randomly more than once, and (iii) structural clustering of SHMs suggests that they act synergistically to improve function, thus SHMs that are part of structural clusters are more likely to be adaptive. To the best of our knowledge, this is a novel approach; while many studies have investigated patterns of convergent SHMs, patterns of *correlations* between convergent SHMs have not been explored. [16, 18, 29]

## Results

### Germline use and identification of SHMs

IMGT’s V-QUEST tool was used to determine the most likely germline V gene for heavy and light chain. [30] A diverse use of germline V genes was found for the antibodies in the data set (Fig. S1, S2, Table S1). On average, heavy-chain germline alleles were used 3.9 times, light-chain alleles 6.5 times, with preferential use of some alleles. With germline V genes assigned, we identified the most likely SHMs for each antibody. A total of 5,468 SHMs were found, 3,493 in the heavy chain and 1,975 in the light chain. 3,435 of the mutations are located in framework regions, 2,033 in CDR loops. On average, each antibody bears 12.0 SHMs, 7.7 in the heavy chain and 4.3 in the light chain. We also compared amino-acid mutabilities between our data set and that of reference, which investigated nucleotide-motif dependent sequence biases for SHMs during a GC reaction. [17] A moderate correlation was found (Fig. S3).

### Convergent SHMs

We next identified convergent SHMs in the data set, i.e. SHMs that occurred in different antibodies but at the same sequence position and with the same germline and mature amino acid identities. A surprisingly large number of such SHMs was found. Out of the 1225 SHMs unique in both sequence position and germline and mature amino acid identity in the heavy chain, 595 occur more than once, and out of the 905 unique somatic mutations in the light chain, 377 occur more than once (Fig. 1). Moreover, most of the convergent SHMs occur at a frequency higher than their respective average, position independent mutation frequency, as can be seen as the enrichment factors in Tables 1, S5, and S6. For example, a 198-fold enrichment was observed for S^H^15P (first entry in Table 1). This mutations occurred in 76.5 % of antibodies in the data set that bear a Ser residue at position 15 in the heavy chain, compared to a 0.4 % frequency of Ser to Pro mutations for all Ser residues in the VH genes of all antibodies. 59.1 % of all S to P mutations in the heavy chain V gene of all antibodies occurred at position 15.

**Figure 1.**
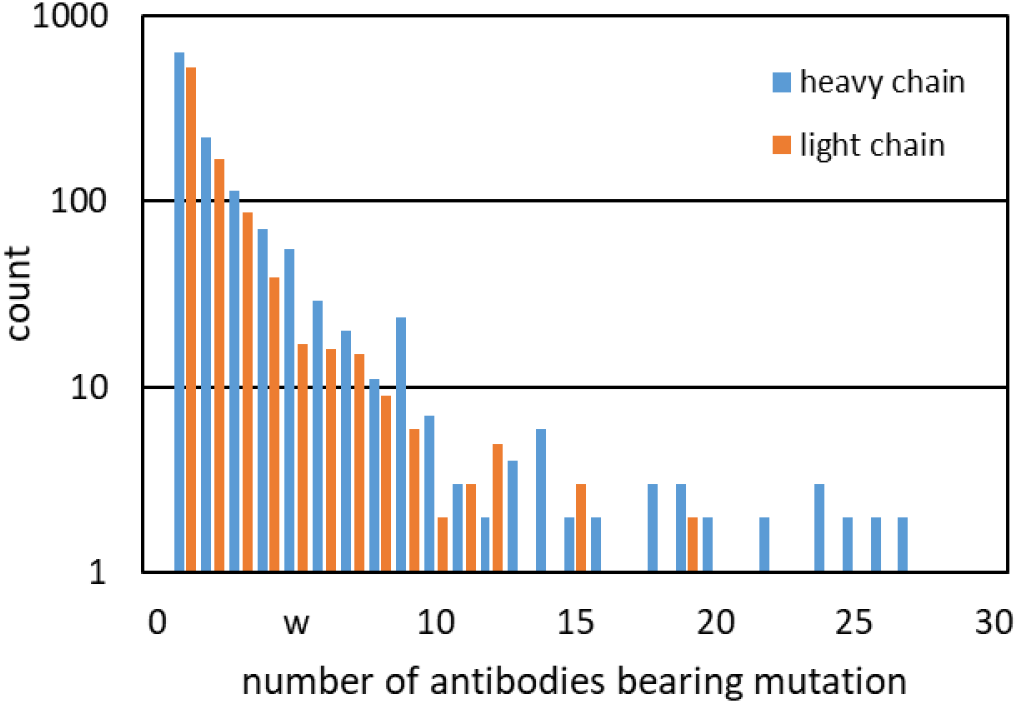
A substantial number of SHMs occurred in more than one antibody (count refers to how many antibodies bear a given SHM).

**Table 1.**
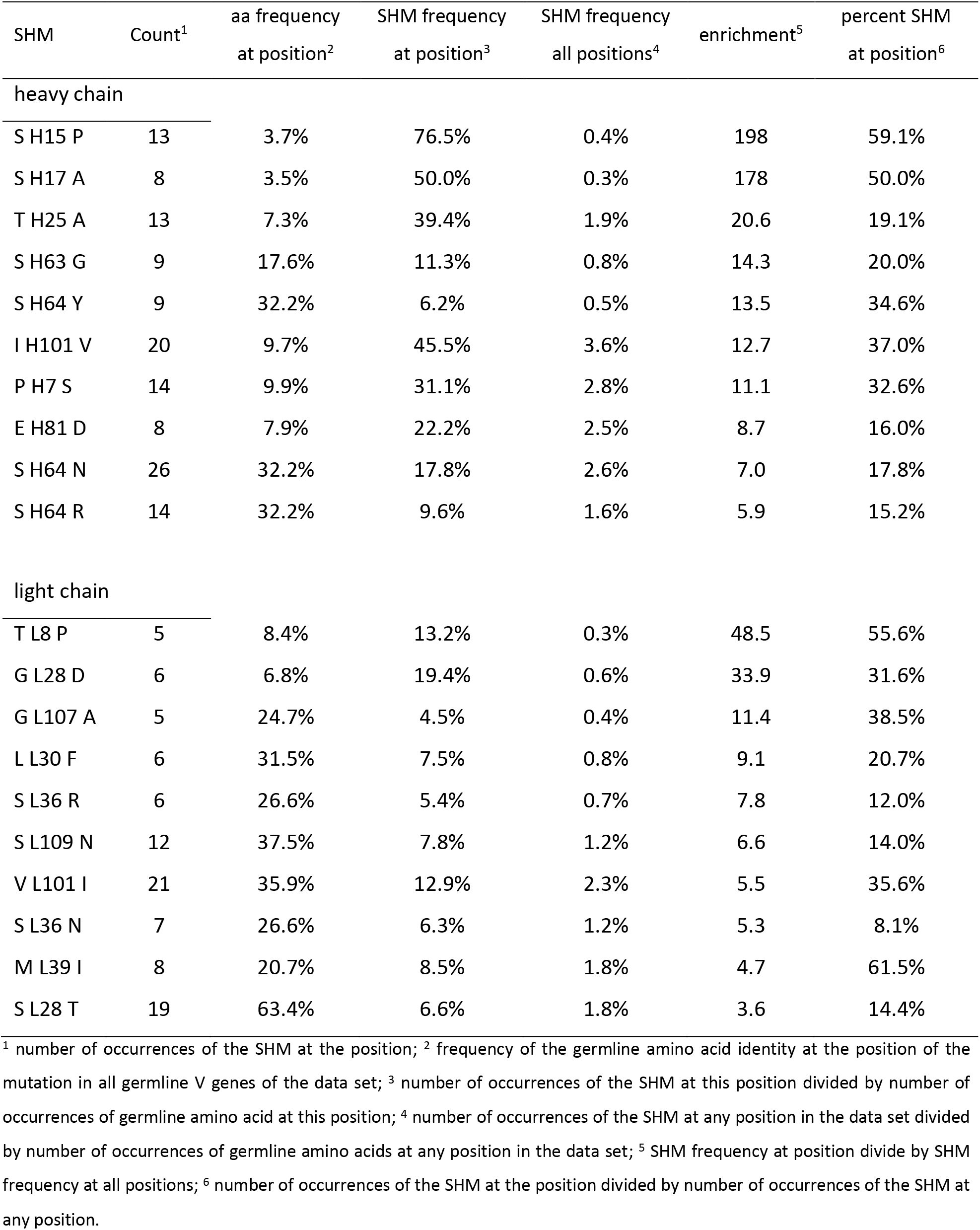
Selected convergent SHMs (for complete list, see Supporting Information).

| SHM | Count<sup>1</sup> | aa frequency<br>at position<sup>2</sup> | SHM frequency<br>at position<sup>3</sup> | SHM frequency<br>all positions<sup>4</sup> | enrichment<sup>5</sup> | percent SHM<br>at position<sup>6</sup> |
| --- | --- | --- | --- | --- | --- | --- |
| heavy chain |  |  |  |  |  |  |
| S H15 P | 13 | 3.7% | 76.5% | 0.4% | 198 | 59.1% |
| S H17 A | 8 | 3.5% | 50.0% | 0.3% | 178 | 50.0% |
| T H25 A | 13 | 7.3% | 39.4% | 1.9% | 20.6 | 19.1% |
| S H63 G | 9 | 17.6% | 11.3% | 0.8% | 14.3 | 20.0% |
| S H64 Y | 9 | 32.2% | 6.2% | 0.5% | 13.5 | 34.6% |
| I H101 V | 20 | 9.7% | 45.5% | 3.6% | 12.7 | 37.0% |
| P H7 S | 14 | 9.9% | 31.1% | 2.8% | 11.1 | 32.6% |
| E H81 D | 8 | 7.9% | 22.2% | 2.5% | 8.7 | 16.0% |
| S H64 N | 26 | 32.2% | 17.8% | 2.6% | 7.0 | 17.8% |
| S H64 R | 14 | 32.2% | 9.6% | 1.6% | 5.9 | 15.2% |
| light chain |  |  |  |  |  |  |
| T L8 P | 5 | 8.4% | 13.2% | 0.3% | 48.5 | 55.6% |
| G L28 D | 6 | 6.8% | 19.4% | 0.6% | 33.9 | 31.6% |
| G L107 A | 5 | 24.7% | 4.5% | 0.4% | 11.4 | 38.5% |
| L L30 F | 6 | 31.5% | 7.5% | 0.8% | 9.1 | 20.7% |
| S L36 R | 6 | 26.6% | 5.4% | 0.7% | 7.8 | 12.0% |
| S L109 N | 12 | 37.5% | 7.8% | 1.2% | 6.6 | 14.0% |
| V L101 I | 21 | 35.9% | 12.9% | 2.3% | 5.5 | 35.6% |
| S L36 N | 7 | 26.6% | 6.3% | 1.2% | 5.3 | 8.1% |
| M L39 I | 8 | 20.7% | 8.5% | 1.8% | 4.7 | 61.5% |
| S L28 T | 19 | 63.4% | 6.6% | 1.8% | 3.6 | 14.4% |

<sup>1</sup> number of occurrences of the SHM at the position; <sup>2</sup> frequency of the germline amino acid identity at the position of the mutation in all germline V genes of the data set; <sup>3</sup> number of occurrences of the SHM at this position divided by number of occurrences of germline amino acid at this position; <sup>4</sup> number of occurrences of the SHM at any position in the data set divided by number of occurrences of germline amino acids at any position in the data set; <sup>5</sup> SHM frequency at position divide by SHM frequency at all positions; <sup>6</sup> number of occurrences of the SHM at the position divided by number of occurrences of the SHM at any position.

### Correlated convergent mutations

We next looked for correlated convergent SHMs, i.e. multiple convergent SHMs occurring together in different antibodies. An astonishing number of such correlated convergent mutations were found. 329 of the 454 antibodies share at least two, 161 share at least three, 87 share at least four, and 53 share at least five SHMs. Tables 2 and S8 show high-frequency correlated convergent SHMs.

**Table 2.**
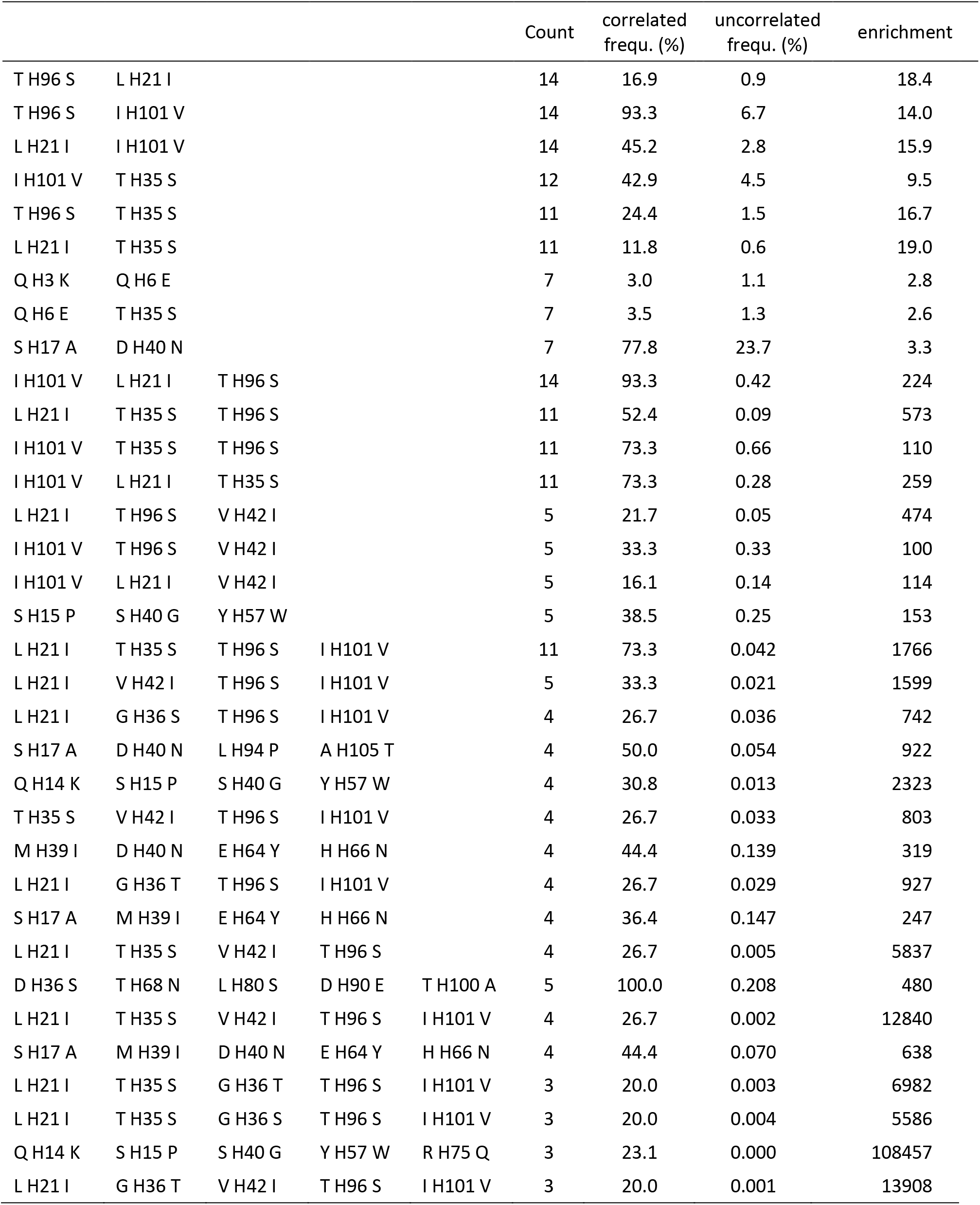
Selected Correlated Convergent Mutations.

As may be expected, enrichment factors, which express the ratio of observed mutation frequency for the group of mutations, i.e. the number of antibodies that bear the mutation divided by the number of antibodies that have the corresponding germline amino acids at the residues in question, to the uncorrelated frequency, i.e. the product of position-dependent SHM frequencies (column 4 in Table 1), increases dramatically with the number of correlated SHMs that occur together in different antibodies. Typical enrichment factors for pairwise correlations are 10-20 range, while enrichment factors for three mutations occurring simultaneously is in the 10^2^ range, those for four or five mutations can be as high as 10^4^ (Table 2, S8).

### Structural clusters of correlated convergent mutations

To further analyze the surprisingly large number of correlated convergent SHMs, we determined pairwise C_α_-C_α_ distances between the correlated mutations (Table S9). We reasoned that short distances may suggest synergistic behavior between the correlated mutations, especially if the SHMs in question are distant in sequence position. 148 instances were found where two convergent SHMs which are shared by several antibodies, are within 12 Å of each other, 13 instances of three, 8 instances of four, and one instance of 5.

Pairwise correlated convergent SHMs were found abundantly within the heavy chain variable region (2392), and to a much lesser extent within the light chain variable region (361) and between heavy and light chain residues (1025, Fig. 2).

**Figure 2.**
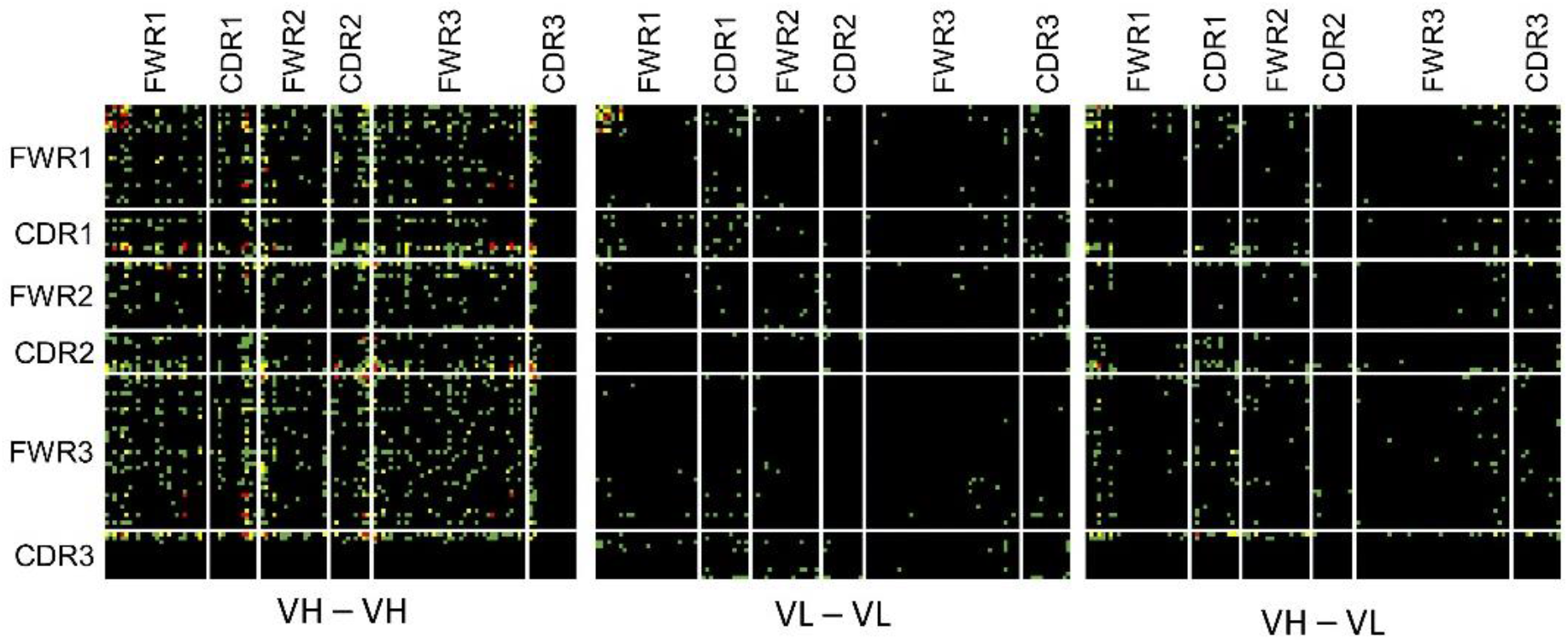
Correlated convergent mutations occur predominantly in the heavy chain. Shown are the sequence locations of pairs of correlated convergent SHMs (left, both mutations are in the heavy chain; middle, both mutations are in the light chain; right, one mutation of the pair is in the heavy, the other in the light chain; coloring: correlated convergent SHM observed in more than 10 (red), 5 (yellow), or less than 5 (green) antibodies of the data set).

## Discussion

Our analysis revealed a surprisingly large number of correlated convergent SHMs. A drawback of computationally assigning germline genes and SHMs is the possibility of error which, if it happens consistently, may result in artefacts. One can easily imagine that a repeatedly misassigned germline may produce a pattern indistinguishable from an actual convergent SHM. In addition, misassigned germlines are more likely to produce high-count clusters of correlated convergent SHMs and without doubt, those are present in our result. Identifying *structural* clusters of convergent SHMs may safeguard against such artefacts since it seems less likely that randomly misassigned SHMs are structurally related in a meaningful way. Therefore, such structural clusters should be prioritized when investigating functional aspects of our results, either computationally using molecular modeling, ore experimentally by introducing or reversing the correlated SHMs in antibody constructs to study their effect on binding affinity, specificity, or thermal stability. Fig. 3 shows one such example of structural clusters of convergent SHMs that are good candidates for follow up studies. Four correlated convergent SHMs were found in four different antibodies, corresponding to PDB structures 1JHL, 1FSK, 4AG4, and 5GIS. [31] [32] [33] [34] The corresponding antibodies are clonally unrelated and each recognize a different antigen, *pheasant* egg white lysozyme (1JHL), *betula pendula* major pollen allergen bet v 1-a (1FSK), *human* epithelial discoidin domaincontaining receptor 1 (4AG4), and *human* isocitrate dehydrogenase 1/2 (5GIS). The four SHMs cluster in two pairs. S^H^17A and L^H^94P have a C_α_-C_α_ distance of 6.6 Å in the crystal structure and are located approximately 30 Å away from the binding site, while D^H^40N and A^H^105T have a C_α_-C_α_ distance of 5.5 Å in the crystal structure and are located approximately 12 Å away from the binding site. In the mature structure, Asn^H^40 and Thr^H^105 form several hydrogen bonds that cross-link the adjacent beta-strands that they are part of. In the germline, the corresponding residues are Asp^H^40 and Ala^H^105, and due to the aliphatic Ala side chain some of the hydrogen bonds cannot be formed. In the other SHM pair, Leu^H^94 is replaced by a Pro^H^94, which rigidifies the loop that this residue is part of. The adjacent Ser^H^17 is replaced by Ala^H^17, perhaps to accommodate the loop structure rigidified by the L^H^94P mutation. Thus, both pairs of SHMs appear to rigidify the framework, a mechanism similar to one we describe previously for antibody 4-4-20. [7] There is no obvious connection between the two pairs of SHMs, so why they would preferentially occur together remains to be seen. Notably, this example, which is just one of the many structural clusters of correlated convergent SHMs we have identified, demonstrates that focusing on this type of mutations may be a suitable strategy to identify patterns of adaptive SHMs that are common between clonally unrelated antibodies.

**Figure 3.**
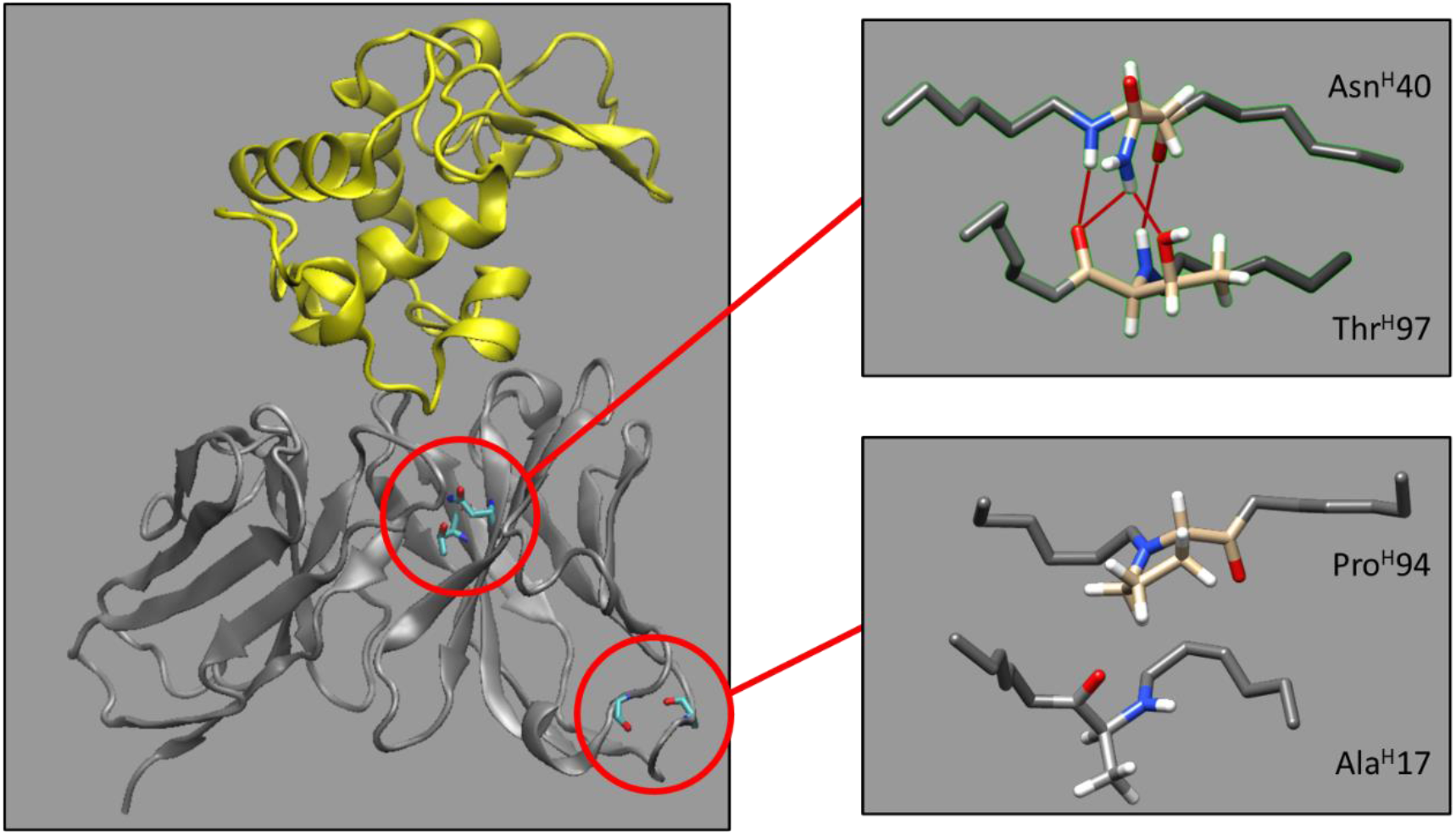
Two concurrently occurring pairs of convergent SHMs may rigidify the framework region. Left: Crystal structure of anti-hen egg lysozyme antibody D11.15 with side chains shown for the two pairs of SHM, and antigen shown in yellow. Right: Enlarged view of the two pairs of SHMs with the backbone of adjacent residues shown in gray and hydrogen bonds shown as bold red lines. PDB ID 1JHL [31].

## Methods

### Creation of a dataset of clonally unrelated PDB structures with gapless germline V gene assignments

We identified 964 murine structures in the dataset created by Sheng et al. which contains 3163 nonredundant, experimentally determined antibody structures from the SAbDab database. [3, 35] We then used IMGT’s DomainGapAlign tool to assign the most likely germline V gene for each antibody. [30]

Only structures with gapless germline alignments for both the VH and VL gene were considered for further analysis, which reduced the number of sequences to 649. We next eliminated highly homologous V gene sequences to exclude clonally related antibodies. This was done by determining the homology of the VH and VL gene for each pair of antibodies and removing structures that were more than 94% homologous in both the VH and VL gene. The 94% cutoff was determined by calculating the number of sequences that would be removed as a function of the cutoff percentage. The first derivative of this function showed a local minimum at 94% (Figure S4), thus we assumed that this was the cutoff between clonally related (with homologies >94%) and clonally unrelated (with homologies ≤94%) sequences. 248 of the 649 sequences were found to be more than 94% homologous in both their VH and VL gene to another sequence. For each pair, the sequence with either the lower number of reported residues in the V regions, or, if those were identical, the sequence with the larger accession number was removed from the data set. This reduced the number of sequences to 525.

To exclude sequences with incorrect germline assignments, we analyzed the distributions of the number of SHMs in the V regions (Figure S5). We reasoned that incorrect germline assignments may lead to an excessive number of SHMs, and therefore removed sequences that had more than 18 mutations in the VH and/or 11 mutations in the VL region. The cutoffs were determined from Gaussian fits to the distribution of the number of SHMs and set at the number of mutations were the Gaussian dropped to 10% of its maximum value. Maximum position (FWHM) of the Gaussian fits were 7.4 (5.7) for VH, 4.0 (3.9) for VL, and 11.3 (6.1) for SHMs of VH and VL combined. 43 structures had more SHMs than the cutoffs in their VH and/or VL region and were removed.

We thus obtained a dataset of 482 PDB structures with VH and VL regions that have sequences for which a gapless germline alignment was found, that are ≤94% mutually homologue in either or both VH and VL and that have not more than 18 assigned SHMs in VH and 11 assigned SHMs in VL. The 482 structures contain 3796 SHMs in VH and 2140 SHMs in VL.

**Supplemental Information** is available on the bioRix website.

