## Supplemental Figures and Tables for "Patterns of convergent somatic hypermutations in the adaptive immune response of Mus musculus"

### Supporting Information

Alexander C. Wenner[1], Charles A. Mettler[1], Ellie M. Sharp, Thomas C. Hansen, Isabella B. Vari, Alexander V. Le, Jörg Zimmermann[2]

Department of Chemistry and Biochemistry, Loyola University Chicago, Chicago, Illinois, USA

[1] These authors contributed equally to the paper.

**Table S1. List of most frequently used heavy-chain and light chain V genes in the data set.**

| heavy-chain V gene[1] | number of usages | light-chain V gene[1] | number of usages |
|---|---|---|---|
| IGHV3-2*02 | 26 | IGKV1-117*01 | 46 |
| IGHV5-6*01 | 26 | IGKV1-110*01 | 29 |
| IGHV1-5*01 | 23 | IGKV10-96*01 | 26 |
| IGHV14-3*02 | 22 | IGKV3-1*01 | 20 |
| IGHV1-7*01 | 17 | IGLV1*01 | 18 |
| IGHV7-3*02 | 15 | IGKV1-135*01 | 16 |
| IGHV1-9*01 | 15 | IGKV8-21*01 | 15 |
| IGHV8-12*01 | 13 | IGKV4-57*01 | 12 |
| IGHV2-6-7*01 | 12 | IGKV12-41*01 | 12 |
| IGHV1-26*01 | 12 | IGKV4-59*01 | 12 |
| IGHV5-12*01 | 12 | IGKV2-137*01 | 11 |
| IGHV5-6-5*01 | 11 | IGKV3-5*01 | 10 |
| IGHV5-9*01 | 11 | IGKV5-43*01 | 9 |
| IGHV9-3*01 | 10 | IGKV8-30*01 | 9 |
| IGHV1-61*01 | 9 | IGKV3-4*01 | 9 |
| IGHV9-2-1*01 | 8 | IGKV3-12*01 | 9 |
| IGHV5-17*02 | 8 | IGKV3-2*01 | 9 |
| IGHV1-4*01 | 8 | IGKV6-20*01 | 9 |
| IGHV5-12-1*01 | 8 | IGKV8-19*01 | 8 |
| IGHV9-3-1*01 | 7 | IGKV12-44*01 | 8 |
| IGHV6-6*01 | 6 | IGKV6-17*01 | 8 |
| IGHV3-8*02 | 6 | IGKV3-10*01 | 8 |
| IGHV2-9*02 | 6 | IGKV14-111*01 | 7 |
| IGHV1-72*01 | 5 | IGKV6-15*01 | 7 |
| IGHV1-15*01 | 5 | IGKV4-55*01 | 7 |
| IGHV1-76*01 | 5 | IGKV13-84*01 | 7 |
| IGHV1-82*01 | 5 | IGKV1-133*01 | 7 |
| IGHV3-6*01 | 5 | IGLV3*01 | 7 |
| IGHV1-69*02 | 5 | IGKV19-93*01 | 6 |
| IGHV8-8*01 | 5 | IGKV14-100*01 | 6 |
| IGHV1S137*01 | 5 | IGKV10-94*01 | 6 |
| IGHV5-6-2*01 | 5 | IGKV5-48*01 | 6 |
| IGHV4-1*02 | 5 | IGKV2-109*01 | 5 |
| IGHV1S136*01 | 5 | IGKV4-57-1*01 | 5 |
| IGHV5-4*02 | 5 | IGKV10-96*02 | 5 |
| IGHV6-6*02 | 5 | IGKV4-74*01 | 5 |
| IGHV1S5*01 | 5 | IGKV3-7*01 | 5 |

[1] in IMGT ontology [Guidicelli and Lefranc (1999), *Bioinformatics* 15:1047]

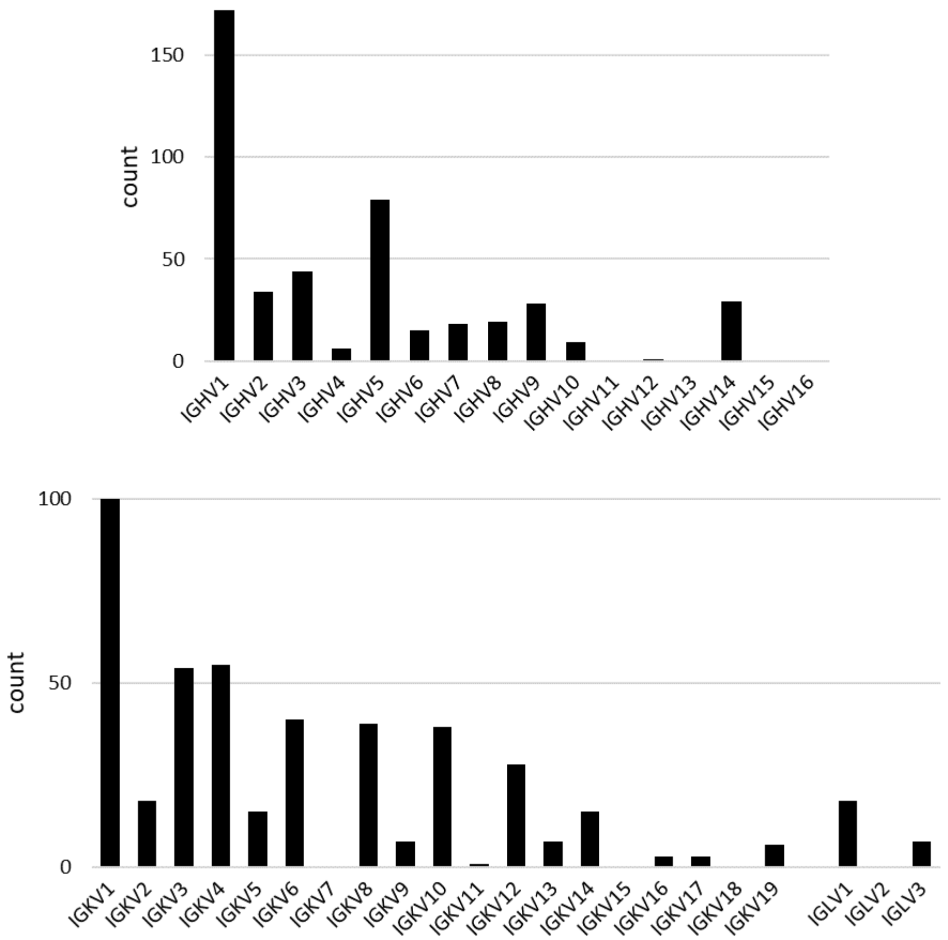

**Figure S1.** Use of germline genes in dataset by IMGT subgroup for heavy chain (top) and light chain (bottom) IGV genes.

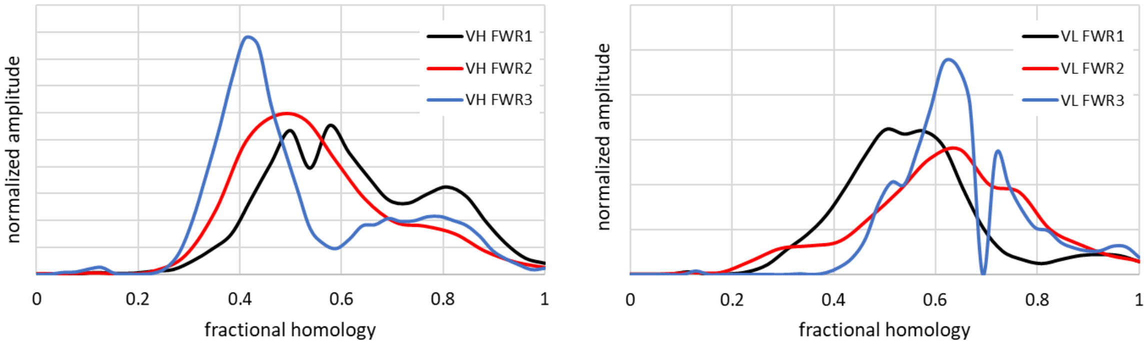

**Figure S2.** Homology distributions for the framework regions of heavy (left) and light chain (right) germline V genes used by the antibodies in the data set.

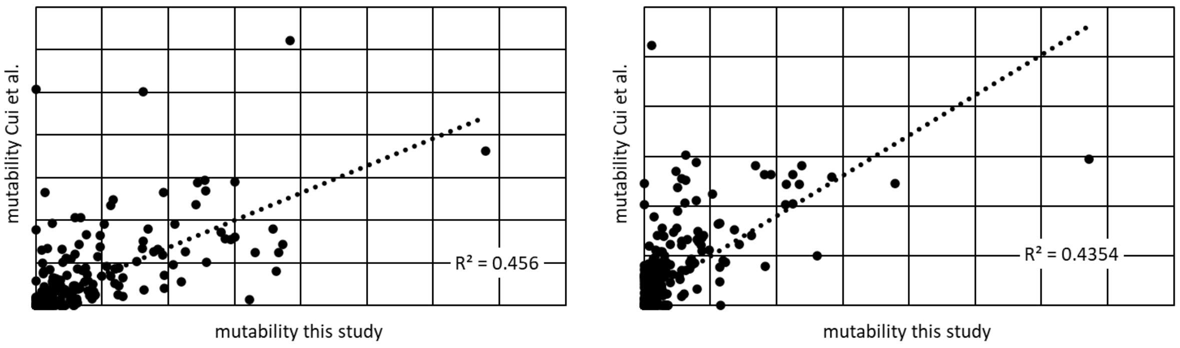

**Figure S3.** Correlation between mutabilities of our data set and Cui et al. (2016) *J. Immunol.* 197:3566 for heavy chain (left) and light chain (right).

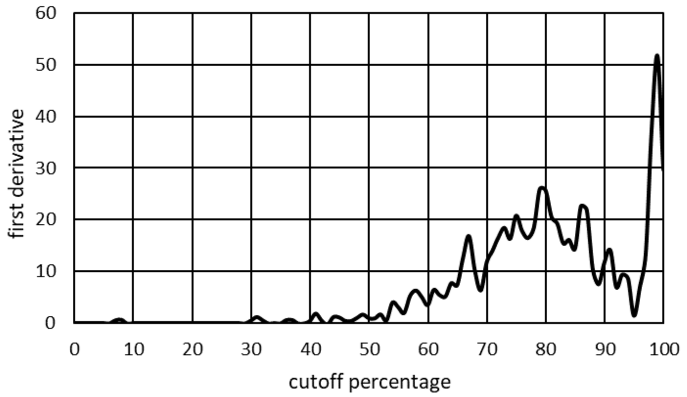

**Figure S4.** First derivative of the number of structures that are less homologous in their VH and/or VL gene as a function of the homology cut-off.

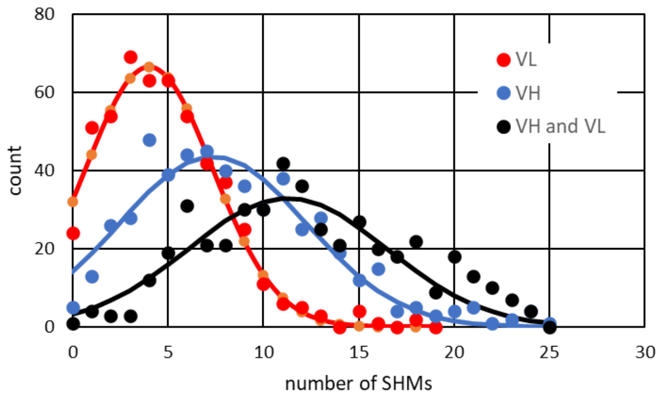

**Figure S5.** Distribution of number of SHM in VH (green), VL (blue) and VH and VL combined (red) of the 525 structures with ≤94% pairwise homology. The circles are actual counts, lines are best fits of a Gaussian functions to the SHM counts.

**Table S2.** Sequence-position independent number of somatic mutations in the heavy chain.

| gl \ ma | A | C | D | E | F | G | H | I | K | L | M | N | P | Q | R | S | T | V | W | Y |
|---|---|---|---|---|---|---|---|---|---|---|---|---|---|---|---|---|---|---|---|---|
| A | 0 | 2 | 5 | 3 | 0 | 36 | 0 | 7 | 0 | 3 | 0 | 1 | 17 | 0 | 2 | 31 | 111 | 87 | 0 | 1 |
| C | 0 | 0 | 0 | 0 | 1 | 0 | 0 | 0 | 0 | 0 | 0 | 0 | 0 | 0 | 0 | 0 | 0 | 0 | 0 | 0 |
| D | 14 | 0 | 0 | 26 | 0 | 17 | 5 | 0 | 5 | 4 | 0 | 36 | 1 | 0 | 0 | 10 | 3 | 11 | 1 | 2 |
| E | 5 | 0 | 50 | 0 | 0 | 7 | 1 | 1 | 15 | 1 | 1 | 2 | 0 | 43 | 3 | 2 | 0 | 5 | 0 | 4 |
| F | 0 | 0 | 0 | 0 | 0 | 0 | 1 | 8 | 0 | 21 | 0 | 0 | 0 | 0 | 0 | 9 | 2 | 7 | 0 | 8 |
| G | 20 | 0 | 72 | 12 | 3 | 0 | 1 | 2 | 2 | 1 | 0 | 8 | 0 | 0 | 15 | 22 | 5 | 14 | 0 | 4 |
| H | 0 | 0 | 1 | 0 | 1 | 0 | 0 | 0 | 0 | 2 | 0 | 9 | 1 | 5 | 7 | 1 | 0 | 0 | 0 | 8 |
| I | 1 | 0 | 1 | 0 | 7 | 1 | 0 | 0 | 2 | 18 | 13 | 3 | 4 | 0 | 0 | 2 | 6 | 54 | 0 | 0 |
| K | 5 | 0 | 2 | 28 | 0 | 2 | 1 | 6 | 0 | 3 | 11 | 34 | 0 | 49 | 110 | 4 | 20 | 6 | 0 | 2 |
| L | 1 | 0 | 1 | 1 | 23 | 0 | 0 | 20 | 0 | 0 | 14 | 0 | 9 | 6 | 1 | 8 | 0 | 30 | 0 | 0 |
| M | 0 | 0 | 0 | 0 | 4 | 0 | 0 | 64 | 2 | 27 | 0 | 0 | 0 | 0 | 0 | 0 | 2 | 16 | 0 | 0 |
| N | 2 | 0 | 36 | 3 | 3 | 3 | 31 | 20 | 39 | 2 | 3 | 0 | 0 | 2 | 7 | 48 | 19 | 1 | 0 | 18 |
| P | 13 | 2 | 0 | 0 | 0 | 0 | 2 | 1 | 0 | 12 | 0 | 1 | 0 | 2 | 2 | 43 | 15 | 1 | 0 | 0 |
| Q | 4 | 0 | 1 | 78 | 0 | 0 | 25 | 0 | 43 | 21 | 2 | 2 | 1 | 0 | 9 | 3 | 1 | 4 | 0 | 0 |
| R | 4 | 4 | 1 | 4 | 1 | 20 | 4 | 10 | 21 | 9 | 1 | 9 | 2 | 9 | 0 | 31 | 21 | 0 | 2 | 7 |
| S | 16 | 2 | 12 | 0 | 13 | 45 | 3 | 19 | 5 | 13 | 2 | 146 | 22 | 0 | 92 | 0 | 118 | 9 | 0 | 26 |
| T | 68 | 0 | 4 | 2 | 1 | 3 | 0 | 86 | 16 | 5 | 4 | 23 | 15 | 1 | 10 | 129 | 0 | 5 | 2 | 1 |
| V | 15 | 0 | 1 | 14 | 5 | 2 | 0 | 39 | 0 | 28 | 18 | 2 | 0 | 10 | 0 | 1 | 0 | 0 | 0 | 1 |
| W | 0 | 1 | 0 | 3 | 1 | 0 | 0 | 0 | 0 | 6 | 0 | 0 | 0 | 0 | 3 | 1 | 1 | 0 | 0 | 2 |
| Y | 1 | 4 | 13 | 4 | 108 | 1 | 28 | 1 | 4 | 5 | 0 | 26 | 0 | 4 | 1 | 35 | 7 | 5 | 11 | 0 |

**Table S3.** Sequence-position independent number of somatic mutations in the light chain.

| gl \ ma | A | C | D | E | F | G | H | I | K | L | M | N | P | Q | R | S | T | V | W | Y |
|---|---|---|---|---|---|---|---|---|---|---|---|---|---|---|---|---|---|---|---|---|
| A | 0 | 0 | 6 | 1 | 0 | 23 | 0 | 2 | 0 | 0 | 0 | 0 | 12 | 0 | 1 | 18 | 45 | 38 | 0 | 0 |
| C | 0 | 0 | 0 | 0 | 0 | 0 | 0 | 0 | 0 | 0 | 0 | 0 | 0 | 0 | 0 | 0 | 0 | 0 | 0 | 0 |
| D | 8 | 0 | 0 | 25 | 0 | 4 | 4 | 2 | 0 | 0 | 0 | 14 | 0 | 1 | 0 | 0 | 2 | 3 | 0 | 7 |
| E | 8 | 0 | 30 | 0 | 0 | 12 | 0 | 1 | 8 | 0 | 0 | 2 | 0 | 2 | 0 | 5 | 0 | 4 | 0 | 1 |
| F | 0 | 1 | 0 | 0 | 0 | 0 | 1 | 2 | 0 | 8 | 0 | 0 | 0 | 0 | 0 | 4 | 0 | 2 | 15 | 3 |
| G | 13 | 0 | 19 | 6 | 0 | 0 | 1 | 1 | 0 | 0 | 0 | 1 | 0 | 1 | 6 | 2 | 1 | 2 | 0 | 0 |
| H | 1 | 0 | 1 | 1 | 0 | 0 | 0 | 0 | 2 | 7 | 0 | 7 | 1 | 4 | 7 | 1 | 0 | 0 | 0 | 16 |
| I | 0 | 0 | 0 | 1 | 6 | 0 | 0 | 0 | 1 | 16 | 10 | 3 | 0 | 0 | 1 | 1 | 5 | 16 | 0 | 0 |
| K | 3 | 0 | 0 | 9 | 0 | 1 | 0 | 1 | 0 | 0 | 5 | 8 | 0 | 7 | 45 | 3 | 13 | 5 | 0 | 1 |
| L | 0 | 0 | 1 | 1 | 29 | 0 | 0 | 9 | 3 | 0 | 20 | 1 | 7 | 4 | 9 | 3 | 0 | 25 | 1 | 4 |
| M | 1 | 0 | 0 | 0 | 2 | 0 | 0 | 13 | 1 | 48 | 0 | 0 | 0 | 0 | 0 | 0 | 0 | 17 | 0 | 0 |
| N | 1 | 0 | 28 | 0 | 2 | 4 | 17 | 10 | 28 | 0 | 0 | 0 | 0 | 0 | 1 | 21 | 14 | 0 | 0 | 18 |
| P | 8 | 0 | 0 | 1 | 8 | 1 | 0 | 2 | 0 | 13 | 1 | 0 | 0 | 1 | 6 | 21 | 4 | 1 | 3 | 6 |
| Q | 1 | 0 | 0 | 19 | 0 | 1 | 19 | 0 | 6 | 7 | 0 | 1 | 3 | 0 | 9 | 3 | 0 | 9 | 0 | 0 |
| R | 3 | 0 | 1 | 1 | 1 | 4 | 3 | 3 | 12 | 1 | 0 | 2 | 1 | 2 | 0 | 5 | 9 | 0 | 1 | 1 |
| S | 13 | 2 | 5 | 2 | 21 | 38 | 4 | 18 | 3 | 3 | 1 | 86 | 8 | 0 | 50 | 0 | 132 | 2 | 1 | 11 |
| T | 68 | 0 | 4 | 2 | 1 | 3 | 0 | 86 | 16 | 5 | 4 | 23 | 15 | 1 | 10 | 129 | 0 | 5 | 2 | 1 |
| V | 15 | 0 | 1 | 14 | 5 | 2 | 0 | 39 | 0 | 28 | 18 | 2 | 0 | 10 | 0 | 1 | 0 | 0 | 0 | 1 |
| W | 0 | 1 | 0 | 3 | 1 | 0 | 0 | 0 | 0 | 6 | 0 | 0 | 0 | 0 | 3 | 1 | 1 | 0 | 0 | 2 |
| Y | 1 | 4 | 13 | 4 | 108 | 1 | 28 | 1 | 4 | 5 | 0 | 26 | 0 | 4 | 1 | 35 | 7 | 5 | 11 | 0 |

**Table S4. Most frequent SHMs in in the V genes of heavy chain and light chain.**

| VH | | | VL | | |
|---|---|---|---|---|---|
| germline identity | mature identity | frequency[1] (%) | germline identity | mature identity | frequency[1] (%) |
| M | I | 6.8 | M | L | 6.7 |
| A | T | 3.8 | Y | F | 3.8 |
| Y | F | 3.7 | K | R | 2.8 |
| K | R | 3.7 | H | Y | 2.6 |
| T | S | 3.6 | M | V | 2.4 |
| I | V | 3.6 | V | I | 2.3 |
| H | N | 3.3 | N | K | 2.2 |
| Q | E | 3.2 | N | D | 2.2 |
| N | S | 3.0 | E | D | 2.1 |
| A | V | 3.0 | T | S | 2.1 |
| H | Y | 2.9 | A | T | 1.9 |
| M | L | 2.9 | S | T | 1.8 |
| P | S | 2.8 | M | I | 1.8 |
| H | R | 2.6 | N | S | 1.7 |
| S | N | 2.6 | A | V | 1.6 |
| E | D | 2.5 | N | Y | 1.4 |
| N | K | 2.4 | N | H | 1.4 |
| T | I | 2.4 | D | E | 1.2 |
| N | D | 2.3 | S | N | 1.2 |
| E | Q | 2.2 | F | W | 1.2 |
| D | N | 2.1 | H | N | 1.1 |
| S | T | 2.1 | H | R | 1.1 |
| N | H | 1.9 | H | L | 1.1 |
| R | S | 1.9 | N | T | 1.1 |
| G | D | 1.9 | V | L | 1.0 |
| T | A | 1.9 | A | G | 1.0 |
| H | Q | 1.8 | Y | H | 0.9 |
| Q | K | 1.8 | P | S | 0.9 |
| M | V | 1.7 | T | I | 0.9 |
| K | Q | 1.6 | E | G | 0.9 |
| F | L | 1.6 | L | F | 0.8 |
| V | I | 1.6 | K | T | 0.8 |
| S | R | 1.6 | N | I | 0.8 |
| D | E | 1.5 | R | K | 0.8 |
| R | T | 1.3 | I | V | 0.8 |
| R | K | 1.3 | I | L | 0.8 |
| N | I | 1.3 | A | S | 0.8 |
| R | G | 1.2 | L | V | 0.7 |

[1] calculated as ratio of number of mutations to number of amino acids with germline identity at any position in the germline sequences of all antibodies in the data set

**Table S5.** Position-dependent SHMs in the heavy chain V genes.

| SHM | Count[1] | aa frequency at position[2] | SHM frequency at position[3] | SHM frequency all positions[4] | enrichment[5] | percent SHM at position[6] |
|---|---|---|---|---|---|---|
| H-A105T | 48 | 88.6% | 12.4% | 3.8% | 3.2 | 43.2% |
| H-M39I | 38 | 66.5% | 12.6% | 6.8% | 1.9 | 59.4% |
| H-Q6E | 32 | 52.2% | 13.5% | 3.2% | 4.2 | 41.0% |
| H-A105V | 30 | 88.6% | 7.8% | 3.0% | 2.6 | 34.5% |
| H-Q3K | 27 | 76.0% | 7.8% | 1.8% | 4.4 | 62.8% |
| H-Y103F | 27 | 85.7% | 6.9% | 3.7% | 1.9 | 25.0% |
| H-S64N | 26 | 32.2% | 17.8% | 2.6% | 7.0 | 17.8% |
| H-T29S | 26 | 63.2% | 9.1% | 3.6% | 2.5 | 20.2% |
| H-S85N | 25 | 57.5% | 9.6% | 2.6% | 3.7 | 17.1% |
| H-T35S | 25 | 55.4% | 10.0% | 3.6% | 2.7 | 19.4% |
| H-S36N | 24 | 39.3% | 13.5% | 2.6% | 5.3 | 16.4% |
| H-G74D | 24 | 64.5% | 8.2% | 1.9% | 4.2 | 33.3% |
| H-T65I | 24 | 81.7% | 6.5% | 2.4% | 2.7 | 27.9% |
| H-Q1E | 22 | 44.4% | 10.9% | 3.2% | 3.4 | 28.2% |
| H-Y37F | 22 | 82.6% | 5.9% | 3.7% | 1.6 | 20.4% |
| H-I101V | 20 | 9.7% | 45.5% | 3.6% | 12.7 | 37.0% |
| H-K72R | 20 | 88.8% | 5.0% | 3.7% | 1.4 | 18.2% |
| H-S36T | 19 | 39.3% | 10.7% | 2.1% | 5.2 | 16.1% |
| H-G63D | 19 | 67.0% | 6.3% | 1.9% | 3.2 | 26.4% |
| H-V42I | 19 | 83.7% | 5.0% | 1.6% | 3.1 | 48.7% |
| H-L21I | 18 | 63.4% | 6.3% | 0.6% | 10.6 | 90.0% |
| H-R106T | 18 | 96.7% | 5.1% | 1.3% | 3.9 | 85.7% |
| H-R106S | 18 | 96.7% | 5.1% | 1.9% | 2.6 | 58.1% |
| H-T96S | 16 | 24.0% | 14.7% | 3.6% | 4.0 | 12.4% |
| H-Y88F | 16 | 77.1% | 4.6% | 3.7% | 1.2 | 14.8% |
| H-S64T | 15 | 32.2% | 10.3% | 2.1% | 5.0 | 12.7% |
| H-K14R | 15 | 59.5% | 5.6% | 3.7% | 1.5 | 13.6% |
| H-K5Q | 14 | 13.0% | 23.7% | 1.6% | 14.5 | 28.6% |
| H-P7S | 14 | 9.9% | 31.1% | 2.8% | 11.1 | 32.6% |
| H-S64R | 14 | 32.2% | 9.6% | 1.6% | 5.9 | 15.2% |
| H-E1Q | 14 | 43.9% | 7.0% | 2.2% | 3.2 | 32.6% |
| H-E6Q | 14 | 47.6% | 6.5% | 2.2% | 3.0 | 32.6% |
| H-T65S | 14 | 81.7% | 3.8% | 3.6% | 1.0 | 10.9% |
| H-S15P | 13 | 3.7% | 76.5% | 0.4% | 198.2 | 59.1% |
| H-T25A | 13 | 7.3% | 39.4% | 1.9% | 20.6 | 19.1% |
| H-T29I | 13 | 63.2% | 4.5% | 2.4% | 1.9 | 15.1% |
| H-K20R | 13 | 71.8% | 4.0% | 3.7% | 1.1 | 11.8% |
| H-S36R | 12 | 39.3% | 6.7% | 1.6% | 4.2 | 13.0% |
| H-A25T | 12 | 69.6% | 3.8% | 3.8% | 1.0 | 10.8% |
| H-S66T | 11 | 15.6% | 15.5% | 2.1% | 7.5 | 9.3% |
| H-N40H | 11 | 28.0% | 8.7% | 1.9% | 4.5 | 35.5% |

Table S5 continued

| SHM | Count[1] | aa frequency at position[2] | SHM frequency at position[3] | SHM frequency all positions[4] | enrichment[5] | percent SHM at position[6] |
|---|---|---|---|---|---|---|
| H-T35I | 11 | 55.4% | 4.4% | 2.4% | 1.8 | 12.8% |
| H-V5Q | 10 | 29.3% | 7.5% | 0.4% | 18.1 | 100.0% |
| H-I2V | 10 | 8.4% | 26.3% | 3.6% | 7.3 | 18.5% |
| H-P68S | 10 | 14.8% | 14.9% | 2.8% | 5.3 | 23.3% |
| H-A105S | 10 | 88.6% | 2.6% | 1.1% | 2.4 | 32.3% |
| H-M101I | 10 | 18.9% | 11.6% | 6.8% | 1.7 | 15.6% |
| H-S92N | 10 | 51.3% | 4.3% | 2.6% | 1.7 | 6.8% |
| H-K70R | 10 | 44.3% | 5.0% | 3.7% | 1.4 | 9.1% |
| H-D40N | 9 | 4.2% | 47.4% | 2.1% | 22.6 | 25.0% |
| H-S63G | 9 | 17.6% | 11.3% | 0.8% | 14.3 | 20.0% |
| H-S64Y | 9 | 32.2% | 6.2% | 0.5% | 13.5 | 34.6% |
| H-A9P | 9 | 27.8% | 7.1% | 0.6% | 12.2 | 52.9% |
| H-D36S | 9 | 36.9% | 5.4% | 0.6% | 9.2 | 90.0% |
| H-S66R | 9 | 15.6% | 12.7% | 1.6% | 7.9 | 9.8% |
| H-Y66H | 9 | 26.7% | 7.4% | 1.0% | 7.7 | 32.1% |
| H-T105S | 9 | 7.6% | 27.3% | 3.6% | 7.5 | 7.0% |
| H-S66N | 9 | 15.6% | 12.7% | 2.6% | 5.0 | 6.2% |
| H-Q5L | 9 | 53.1% | 3.7% | 0.9% | 4.3 | 42.9% |
| H-R106I | 9 | 96.7% | 2.5% | 0.6% | 4.1 | 90.0% |
| H-K70N | 9 | 44.3% | 4.5% | 1.1% | 3.9 | 26.5% |
| H-S57N | 9 | 25.8% | 7.7% | 2.6% | 3.0 | 6.2% |
| H-L50F | 9 | 98.9% | 2.0% | 0.7% | 3.0 | 39.1% |
| H-Y37N | 9 | 82.6% | 2.4% | 0.9% | 2.7 | 34.6% |
| H-Q90H | 9 | 72.5% | 2.7% | 1.0% | 2.6 | 36.0% |
| H-L12V | 9 | 95.6% | 2.1% | 0.9% | 2.3 | 30.0% |
| H-R106G | 9 | 96.7% | 2.5% | 1.2% | 2.0 | 45.0% |
| H-A105G | 9 | 88.6% | 2.3% | 1.2% | 1.9 | 25.0% |
| H-S85T | 9 | 57.5% | 3.4% | 2.1% | 1.7 | 7.6% |
| H-A87V | 9 | 50.0% | 4.0% | 3.0% | 1.3 | 10.3% |
| H-A17T | 9 | 39.2% | 5.1% | 3.8% | 1.3 | 8.1% |
| H-K75R | 9 | 45.0% | 4.4% | 3.7% | 1.2 | 8.2% |
| H-I56V | 9 | 99.1% | 2.0% | 3.6% | 0.6 | 16.7% |
| H-S17A | 8 | 3.5% | 50.0% | 0.3% | 178.2 | 50.0% |
| H-E81D | 8 | 7.9% | 22.2% | 2.5% | 8.7 | 16.0% |
| H-S62T | 8 | 22.9% | 10.1% | 2.1% | 4.9 | 6.8% |
| H-V42M | 8 | 83.7% | 2.1% | 0.7% | 2.8 | 44.4% |
| H-N40S | 8 | 28.0% | 6.3% | 3.0% | 2.1 | 16.7% |
| H-S79T | 8 | 45.6% | 3.9% | 2.1% | 1.9 | 6.8% |
| H-A100G | 8 | 95.4% | 1.8% | 1.2% | 1.5 | 22.2% |
| H-T65A | 8 | 81.7% | 2.2% | 1.9% | 1.1 | 11.8% |
| H-N68S | 8 | 57.3% | 3.1% | 3.0% | 1.0 | 16.7% |
| H-T99S | 8 | 57.9% | 3.0% | 3.6% | 0.8 | 6.2% |

Table S5 continued

| SHM | Count[1] | aa frequency at position[2] | SHM frequency at position[3] | SHM frequency all positions[4] | enrichment[5] | percent SHM at position[6] |
|---|---|---|---|---|---|---|
| H-M89I | 8 | 37.2% | 4.7% | 6.8% | 0.7 | 12.5% |
| H-S64I | 7 | 32.2% | 4.8% | 0.3% | 14.4 | 36.8% |
| H-K66E | 7 | 11.9% | 13.0% | 0.9% | 13.9 | 25.0% |
| H-Y57N | 7 | 18.9% | 8.1% | 0.9% | 9.1 | 26.9% |
| H-V5E | 7 | 29.3% | 5.3% | 0.6% | 9.0 | 50.0% |
| H-D1E | 7 | 11.7% | 13.2% | 1.5% | 8.7 | 26.9% |
| H-L94P | 7 | 83.5% | 1.8% | 0.3% | 7.0 | 77.8% |
| H-N93S | 7 | 11.0% | 14.0% | 3.0% | 4.7 | 14.6% |
| H-S57T | 7 | 25.8% | 6.0% | 2.1% | 2.9 | 5.9% |
| H-V101I | 7 | 39.9% | 3.9% | 1.6% | 2.4 | 17.9% |
| H-N92K | 7 | 35.2% | 4.4% | 2.4% | 1.8 | 17.9% |
| H-Y38F | 7 | 24.4% | 6.3% | 3.7% | 1.7 | 6.5% |
| H-K82R | 7 | 25.3% | 6.1% | 3.7% | 1.7 | 6.4% |
| H-K84R | 7 | 37.0% | 4.2% | 3.7% | 1.1 | 6.4% |
| H-Q5E | 7 | 53.1% | 2.9% | 3.2% | 0.9 | 9.0% |
| H-T77I | 7 | 71.4% | 2.2% | 2.4% | 0.9 | 8.1% |
| H-K43R | 7 | 50.0% | 3.1% | 3.7% | 0.8 | 6.4% |
| H-A25V | 7 | 69.6% | 2.2% | 3.0% | 0.7 | 8.0% |
| H-E97D | 7 | 86.6% | 1.8% | 2.5% | 0.7 | 14.0% |
| H-T86S | 7 | 64.3% | 2.4% | 3.6% | 0.7 | 5.4% |
| H-E51D | 7 | 93.6% | 1.6% | 2.5% | 0.6 | 14.0% |
| H-K5V | 6 | 13.0% | 10.2% | 0.2% | 50.8 | 100.0% |
| H-Y38T | 6 | 24.4% | 5.4% | 0.2% | 22.4 | 85.7% |
| H-Y57W | 6 | 18.9% | 7.0% | 0.4% | 18.4 | 54.5% |
| H-V80A | 6 | 21.6% | 6.1% | 0.6% | 9.8 | 40.0% |
| H-N61K | 6 | 83.3% | 17.1% | 2.4% | 7.0 | 15.4% |
| H-D59N | 6 | 9.5% | 14.0% | 2.1% | 6.6 | 16.7% |
| H-G64D | 6 | 11.0% | 12.0% | 1.9% | 6.2 | 8.3% |
| H-I21M | 6 | 25.1% | 5.3% | 0.9% | 6.1 | 46.2% |
| H-G63V | 6 | 67.0% | 2.0% | 0.4% | 5.3 | 42.9% |
| H-K3Q | 6 | 17.6% | 7.5% | 1.6% | 4.6 | 12.2% |
| H-G54A | 6 | 71.4% | 1.9% | 0.5% | 3.5 | 30.0% |
| H-P45S | 6 | 14.1% | 9.4% | 2.8% | 3.4 | 14.0% |
| H-M101L | 6 | 18.9% | 7.0% | 2.9% | 2.4 | 22.2% |
| H-Y57F | 6 | 18.9% | 7.0% | 3.7% | 1.9 | 5.6% |
| H-R75K | 6 | 55.0% | 2.4% | 1.3% | 1.8 | 28.6% |
| H-K72E | 6 | 88.8% | 1.5% | 0.9% | 1.6 | 21.4% |
| H-P46T | 6 | 86.6% | 1.5% | 1.0% | 1.6 | 40.0% |
| H-Q44L | 6 | 97.6% | 1.4% | 0.9% | 1.6 | 28.6% |
| H-K48Q | 6 | 55.9% | 2.4% | 1.6% | 1.4 | 12.2% |
| H-S85R | 6 | 57.5% | 2.3% | 1.6% | 1.4 | 6.5% |
| H-Y88S | 6 | 77.1% | 1.7% | 1.2% | 1.4 | 17.1% |

Table S5 continued

| SHM | Count[1] | aa frequency at position[2] | SHM frequency at position[3] | SHM frequency all positions[4] | enrichment[5] | percent SHM at position[6] |
|---|---|---|---|---|---|---|
| H-H40Y | 6 | 32.8% | 4.0% | 2.9% | 1.4 | 75.0% |
| H-A9T | 6 | 27.8% | 4.8% | 3.8% | 1.2 | 5.4% |
| H-M39V | 6 | 66.5% | 2.0% | 1.7% | 1.2 | 37.5% |
| H-F30L | 6 | 72.0% | 1.8% | 1.6% | 1.1 | 28.6% |
| H-M39L | 6 | 66.5% | 2.0% | 2.9% | 0.7 | 22.2% |
| H-I78V | 6 | 54.6% | 2.4% | 3.6% | 0.7 | 11.1% |
| H-Y67F | 6 | 99.6% | 1.3% | 3.7% | 0.4 | 5.6% |
| H-Y102F | 6 | 99.8% | 1.3% | 3.7% | 0.4 | 5.6% |
| H-P94L | 5 | 1.1% | 100.0% | 0.8% | 128.2 | 41.7% |
| H-L80S | 5 | 7.3% | 15.2% | 0.2% | 64.1 | 62.5% |
| H-T68N | 5 | 3.1% | 35.7% | 0.6% | 55.1 | 21.7% |
| H-T100A | 5 | 1.1% | 100.0% | 1.9% | 52.2 | 7.4% |
| H-D90E | 5 | 1.5% | 71.4% | 1.5% | 47.1 | 19.2% |
| H-S86R | 5 | 3.3% | 33.3% | 1.6% | 20.7 | 5.4% |
| H-G36S | 5 | 12.8% | 8.6% | 0.6% | 14.6 | 22.7% |
| H-N59Y | 5 | 9.5% | 11.6% | 1.1% | 10.3 | 27.8% |
| H-Y66D | 5 | 26.7% | 4.1% | 0.4% | 9.2 | 38.5% |
| H-K66N | 5 | 11.9% | 9.3% | 1.1% | 8.2 | 14.7% |
| H-K82T | 5 | 25.3% | 4.3% | 0.7% | 6.5 | 25.0% |
| H-Y67L | 5 | 99.6% | 1.1% | 0.2% | 6.4 | 100.0% |
| H-T82K | 5 | 38.3% | 2.9% | 0.5% | 6.4 | 31.3% |
| H-V42F | 5 | 83.7% | 1.3% | 0.2% | 6.3 | 100.0% |
| H-S40G | 5 | 23.3% | 4.7% | 0.8% | 6.0 | 11.1% |
| H-K3N | 5 | 17.6% | 6.3% | 1.1% | 5.5 | 14.7% |
| H-D64N | 5 | 11.9% | 9.3% | 2.1% | 4.4 | 13.9% |
| H-A24S | 5 | 25.6% | 4.3% | 1.1% | 4.0 | 16.1% |
| H-M21L | 5 | 10.1% | 10.9% | 2.9% | 3.8 | 18.5% |
| H-S36G | 5 | 39.3% | 2.8% | 0.8% | 3.6 | 11.1% |
| H-A84T | 5 | 8.1% | 13.5% | 3.8% | 3.5 | 4.5% |
| H-N66H | 5 | 16.3% | 6.8% | 1.9% | 3.5 | 16.1% |
| H-S63T | 5 | 17.6% | 6.3% | 2.1% | 3.0 | 4.2% |
| H-D36G | 5 | 36.9% | 3.0% | 1.0% | 3.0 | 29.4% |
| H-Q14K | 5 | 20.9% | 5.3% | 1.8% | 3.0 | 11.6% |
| H-N82I | 5 | 30.2% | 3.6% | 1.3% | 2.9 | 25.0% |
| H-V19M | 5 | 50.7% | 2.2% | 0.7% | 2.9 | 27.8% |
| H-N66K | 5 | 16.3% | 6.8% | 2.4% | 2.8 | 12.8% |
| H-S74R | 5 | 25.8% | 4.3% | 1.6% | 2.6 | 5.4% |
| H-S58N | 5 | 16.3% | 6.8% | 2.6% | 2.6 | 3.4% |
| H-N57D | 5 | 18.7% | 5.9% | 2.3% | 2.6 | 13.9% |
| H-P58T | 5 | 43.6% | 2.5% | 1.0% | 2.6 | 33.3% |
| H-A68V | 5 | 14.5% | 7.6% | 3.0% | 2.5 | 5.7% |
| H-T96A | 5 | 24.0% | 4.6% | 1.9% | 2.4 | 7.4% |

Table S5 continued

| SHM | Count[1] | aa frequency at position[2] | SHM frequency at position[3] | SHM frequency all positions[4] | enrichment[5] | percent SHM at position[6] |
|---|---|---|---|---|---|---|
| H-L91V | 5 | 54.4% | 2.0% | 0.9% | 2.3 | 16.7% |
| H-D36E | 5 | 36.9% | 3.0% | 1.5% | 2.0 | 19.2% |
| H-S35R | 5 | 35.1% | 3.1% | 1.6% | 1.9 | 5.4% |
| H-K95R | 5 | 17.2% | 6.4% | 3.7% | 1.7 | 4.5% |
| H-S74N | 5 | 25.8% | 4.3% | 2.6% | 1.7 | 3.4% |
| H-N68T | 5 | 57.3% | 1.9% | 1.2% | 1.6 | 26.3% |
| H-S35T | 5 | 35.1% | 3.1% | 2.1% | 1.5 | 4.2% |
| H-N62S | 5 | 32.5% | 4.5% | 3.0% | 1.5 | 10.4% |
| H-P15A | 5 | 94.7% | 1.2% | 0.8% | 1.4 | 38.5% |
| H-S92R | 5 | 51.3% | 2.1% | 1.6% | 1.3 | 5.4% |
| H-G11D | 5 | 44.5% | 2.5% | 1.9% | 1.3 | 6.9% |
| H-L4V | 5 | 99.6% | 1.1% | 0.9% | 1.2 | 16.7% |
| H-S35N | 5 | 35.1% | 3.1% | 2.6% | 1.2 | 3.4% |
| H-V42L | 5 | 83.7% | 1.3% | 1.2% | 1.1 | 17.9% |
| H-S92T | 5 | 51.3% | 2.1% | 2.1% | 1.0 | 4.2% |
| H-H40N | 5 | 32.8% | 3.4% | 3.3% | 1.0 | 55.6% |
| H-Y67S | 5 | 99.6% | 1.1% | 1.2% | 0.9 | 14.3% |
| H-K72Q | 5 | 88.8% | 1.2% | 1.6% | 0.8 | 10.2% |
| H-T86I | 5 | 64.3% | 1.7% | 2.4% | 0.7 | 5.8% |
| H-A76T | 5 | 43.6% | 2.5% | 3.8% | 0.7 | 4.5% |
| H-S93N | 5 | 80.8% | 1.4% | 2.6% | 0.5 | 3.4% |
| H-Q90E | 5 | 72.5% | 1.5% | 3.2% | 0.5 | 6.4% |

[1] number of occurrences of the SHM at the position

[2] frequency of the germline amino acid identity at the position of the mutation in all germline V genes of the data set

[3] number of occurrences of the SHM at this position divided by number of occurrences of germline amino acid at this position

[4] number of occurrences of the SHM at any position in the data set divided by number of occurrences of germline amino acids at any position in the data set

[5] SHM frequency at position divide by SHM frequency at all positions

[6] number of occurrences of the SHM at the position divided by number of occurrences of the SHM at any position

**Table S6.** Position-dependent SHMs in the light chain V genes.

| SHM | Count[1] | aa frequency at position[2] | SHM frequency at position[3] | SHM frequency all positions[4] | enrichment[5] | percent SHM at position[6] |
|---|---|---|---|---|---|---|
| L-M4L | 29 | 65.4% | 9.8% | 6.7% | 1.5 | 60.4% |
| L-T7S | 22 | 31.3% | 15.5% | 2.1% | 7.2 | 31.0% |
| L-V101I | 21 | 35.9% | 12.9% | 2.3% | 5.5 | 35.6% |
| L-S28T | 19 | 63.4% | 6.6% | 1.8% | 3.6 | 14.4% |
| L-Y38F | 19 | 75.1% | 5.6% | 3.8% | 1.5 | 19.6% |
| L-F116W | 15 | 34.0% | 93.8% | 1.2% | 81.1 | 100.0% |
| L-S92N | 15 | 72.7% | 4.5% | 1.2% | 3.8 | 17.4% |
| L-Y42F | 15 | 80.0% | 4.1% | 3.8% | 1.1 | 15.5% |
| L-D1E | 13 | 76.9% | 3.7% | 1.2% | 3.0 | 52.0% |
| L-L4M | 12 | 28.9% | 9.2% | 0.6% | 16.0 | 60.0% |
| L-P116L | 12 | 51.1% | 50.0% | 0.5% | 92.3 | 92.3% |
| L-S109N | 12 | 37.5% | 7.8% | 1.2% | 6.6 | 14.0% |
| L-V2I | 12 | 22.9% | 11.5% | 2.3% | 4.9 | 20.3% |
| L-Y55F | 12 | 86.6% | 3.1% | 3.8% | 0.8 | 12.4% |
| L-S109T | 11 | 37.5% | 7.2% | 1.8% | 3.9 | 8.3% |
| L-S22T | 11 | 60.4% | 4.0% | 1.8% | 2.2 | 8.3% |
| L-Y103F | 11 | 67.8% | 3.6% | 3.8% | 0.9 | 11.3% |
| L-S107T | 10 | 25.8% | 8.5% | 1.8% | 4.7 | 7.6% |
| L-S26T | 10 | 100.0% | 2.2% | 1.8% | 1.2 | 7.6% |
| L-L3V | 9 | 12.8% | 15.5% | 0.7% | 21.7 | 36.0% |
| L-N34D | 9 | 58.8% | 7.3% | 2.2% | 3.2 | 32.1% |
| L-P46S | 9 | 86.1% | 2.3% | 0.9% | 2.6 | 42.9% |
| L-Q3V | 9 | 19.8% | 10.0% | 0.3% | 33.9 | 100.0% |
| L-S7T | 9 | 60.8% | 3.3% | 1.8% | 1.8 | 6.8% |
| L-V3Q | 9 | 64.5% | 3.1% | 0.4% | 8.6 | 100.0% |
| L-I2V | 8 | 69.8% | 2.5% | 0.8% | 3.2 | 50.0% |
| L-K45R | 8 | 88.5% | 2.0% | 2.8% | 0.7 | 17.8% |
| L-K51R | 8 | 78.0% | 2.3% | 2.8% | 0.8 | 17.8% |
| L-M39I | 8 | 20.7% | 8.5% | 1.8% | 4.7 | 61.5% |
| L-Q3E | 8 | 19.8% | 8.9% | 0.6% | 14.3 | 42.1% |
| L-S37N | 8 | 30.0% | 5.9% | 1.2% | 4.9 | 9.3% |
| L-S37T | 8 | 30.0% | 5.9% | 1.8% | 3.2 | 6.1% |
| L-S5T | 8 | 5.9% | 29.6% | 1.8% | 16.2 | 6.1% |
| L-Y38H | 8 | 75.1% | 2.3% | 0.9% | 2.6 | 34.8% |
| L-*116Y | 7 | 0.0% | #DIV/0! | 0.0% | #DIV/0! | #DIV/0! |
| L-A57T | 7 | 43.4% | 3.6% | 1.9% | 1.9 | 15.6% |
| L-A96T | 7 | 56.2% | 2.7% | 1.9% | 1.4 | 15.6% |
| L-E97D | 7 | 92.7% | 1.7% | 2.1% | 0.8 | 23.3% |
| L-M39L | 7 | 20.7% | 7.4% | 6.7% | 1.1 | 14.6% |
| L-N36D | 7 | 28.5% | 5.9% | 2.2% | 2.6 | 25.0% |
| L-N36K | 7 | 28.5% | 5.9% | 2.2% | 2.6 | 25.0% |

Table S7 continued.

| SHM | Count[1] | aa frequency at position[2] | SHM frequency at position[3] | SHM frequency all positions[4] | enrichment[5] | percent SHM at position[6] |
|---|---|---|---|---|---|---|
| L-N66K | 7 | 52.2% | 3.0% | 2.2% | 1.3 | 25.0% |
| L-Q106H | 7 | 83.9% | 1.8% | 0.6% | 2.9 | 36.8% |
| L-S108T | 7 | 24.5% | 6.3% | 1.8% | 3.5 | 5.3% |
| L-S28N | 7 | 63.4% | 2.4% | 1.2% | 2.0 | 8.1% |
| L-S36N | 7 | 26.6% | 6.3% | 1.2% | 5.3 | 8.1% |
| L-V2L | 7 | 22.9% | 6.7% | 1.0% | 6.5 | 26.9% |
| L-Y107F | 7 | 16.7% | 9.2% | 3.8% | 2.4 | 7.2% |
| L-Y55H | 7 | 86.6% | 1.8% | 0.9% | 2.0 | 30.4% |
| L-A57V | 6 | 43.4% | 3.0% | 1.6% | 1.9 | 15.8% |
| L-A74V | 6 | 25.8% | 5.1% | 1.6% | 3.2 | 15.8% |
| L-G28D | 6 | 6.8% | 19.4% | 0.6% | 33.9 | 31.6% |
| L-I2L | 6 | 69.8% | 1.9% | 0.8% | 2.4 | 37.5% |
| L-K51N | 6 | 78.0% | 1.7% | 0.5% | 3.4 | 75.0% |
| L-L116R | 6 | 10.6% | 120.0% | 0.3% | 466.7 | 66.7% |
| L-L30F | 6 | 31.5% | 7.5% | 0.8% | 9.1 | 20.7% |
| L-L52V | 6 | 77.1% | 1.7% | 0.7% | 2.4 | 24.0% |
| L-N1D | 6 | 4.0% | 33.3% | 2.2% | 14.9 | 21.4% |
| L-N40S | 6 | 23.3% | 5.7% | 1.7% | 3.4 | 28.6% |
| L-P116F | 6 | 51.1% | 25.0% | 0.3% | 75.0 | 75.0% |
| L-P116Y | 6 | 51.1% | 25.0% | 0.3% | 100.0 | 100.0% |
| L-S26N | 6 | 100.0% | 1.3% | 1.2% | 1.1 | 7.0% |
| L-S36R | 6 | 26.6% | 5.4% | 0.7% | 7.8 | 12.0% |
| L-T108S | 6 | 11.9% | 11.1% | 2.1% | 5.2 | 8.5% |
| L-Y114F | 6 | 19.1% | 7.1% | 3.8% | 1.9 | 6.2% |
| L-E97A | 5 | 92.7% | 1.2% | 0.6% | 2.1 | 62.5% |
| L-G107A | 5 | 24.7% | 4.5% | 0.4% | 11.4 | 38.5% |
| L-H31Y | 5 | 40.6% | 5.4% | 2.6% | 2.1 | 31.3% |
| L-H40N | 5 | 26.2% | 4.2% | 1.1% | 3.7 | 71.4% |
| L-K24R | 5 | 29.3% | 3.8% | 2.8% | 1.3 | 11.1% |
| L-K90R | 5 | 28.2% | 3.9% | 2.8% | 1.4 | 11.1% |
| L-L52F | 5 | 77.1% | 1.4% | 0.8% | 1.7 | 17.2% |
| L-P116R | 5 | 51.1% | 20.8% | 0.3% | 83.3 | 83.3% |
| L-Q105H | 5 | 53.3% | 2.1% | 0.6% | 3.3 | 26.3% |
| L-S77R | 5 | 77.5% | 1.4% | 0.7% | 2.1 | 10.0% |
| L-S79R | 5 | 100.0% | 1.1% | 0.7% | 1.6 | 10.0% |
| L-T37S | 5 | 35.9% | 3.1% | 2.1% | 1.4 | 7.0% |
| L-T7I | 5 | 31.3% | 3.5% | 0.9% | 4.0 | 17.2% |
| L-T8P | 5 | 8.4% | 13.2% | 0.3% | 48.5 | 55.6% |
| L-V3E | 5 | 64.5% | 1.7% | 0.2% | 8.6 | 100.0% |
| L-Y102F | 5 | 100.0% | 1.1% | 3.8% | 0.3 | 5.2% |
| L-Y87F | 5 | 32.4% | 3.4% | 3.8% | 0.9 | 5.2% |

[1] number of occurrences of the SHM at the position
[2] frequency of the germline amino acid identity at the position of the mutation in all germline V genes of the data set
[3] number of occurrences of the SHM at this position divided by number of occurrences of germline amino acid at this position
[4] number of occurrences of the SHM at any position in the data set divided by number of occurrences of germline amino acids at any position in the data set
[5] SHM frequency at position divide by SHM frequency at all positions
[6] number of occurrences of the SHM at the position divided by number of occurrences of the SHM at any position

**Table S8.** Correlated Convergent Mutations

| | | Count | correlated frequ. (%) | uncorrelated frequ. (%) | enrichment |
|---|---|---|---|---|---|
| T H96 S | L H21 I | 14 | 16.9 | 0.9 | 18.4 |
| T H96 S | I H101 V | 14 | 93.3 | 6.7 | 14.0 |
| L H21 I | I H101 V | 14 | 45.2 | 2.8 | 15.9 |
| I H101 V | T H35 S | 12 | 42.9 | 4.5 | 9.5 |
| T H96 S | T H35 S | 11 | 24.4 | 1.5 | 16.7 |
| L H21 I | T H35 S | 11 | 11.8 | 0.6 | 19.0 |
| Q H3 K | Q H6 E | 7 | 3.0 | 1.1 | 2.8 |
| Q H6 E | T H35 S | 7 | 3.5 | 1.3 | 2.6 |
| S H17 A | D H40 N | 7 | 77.8 | 23.7 | 3.3 |
| A H105 T | M H39 I | 6 | 2.4 | 1.6 | 1.6 |
| S H36 T | Q H6 E | 6 | 6.3 | 1.4 | 4.4 |
| D H40 N | L H94 P | 6 | 31.6 | 0.9 | 36.1 |
| A H25 T | N H40 H | 6 | 8.3 | 0.3 | 25.3 |
| V L2 L | D L1 E | 6 | 5.8 | 0.3 | 23.0 |
| M L4 L | A H105 V | 5 | 2.0 | 0.8 | 2.6 |
| M L4 L | E H1 Q | 5 | 3.7 | 0.7 | 5.4 |
| M L4 L | Q L3 E | 5 | 5.7 | 0.9 | 6.6 |
| T H96 S | V H42 I | 5 | 11.1 | 0.7 | 15.1 |
| Q H1 E | Q H3 K | 5 | 2.8 | 0.9 | 3.3 |
| Q H1 E | Q H6 E | 5 | 3.2 | 1.5 | 2.2 |
| V H42 I | Q H6 E | 5 | 2.1 | 0.7 | 3.2 |
| V H42 I | L H21 I | 5 | 2.3 | 0.3 | 7.3 |
| V H42 I | I H101 V | 5 | 11.4 | 2.3 | 5.0 |
| V H42 I | T H25 A | 5 | 15.2 | 2.0 | 7.7 |
| A H105 T | S H17 A | 5 | 33.3 | 6.2 | 5.4 |
| A H105 T | N H40 H | 5 | 4.5 | 1.1 | 4.2 |
| M H39 I | Q H6 E | 5 | 2.8 | 1.7 | 1.7 |
| M H39 I | T H35 S | 5 | 3.2 | 1.3 | 2.5 |
| M H39 I | Y H88 F | 5 | 1.7 | 0.6 | 3.0 |
| S H64 N | T H35 S | 5 | 7.0 | 1.8 | 4.0 |
| S H85 N | Q H6 E | 5 | 2.6 | 1.3 | 2.0 |
| Q H6 E | I H101 V | 5 | 17.9 | 6.1 | 2.9 |
| Q H6 E | T L7 S | 5 | 7.2 | 2.1 | 3.5 |
| Q H6 E | Q H5 L | 5 | 2.6 | 0.5 | 5.1 |
| T H35 S | T H29 S | 5 | 2.6 | 0.9 | 2.8 |
| T H25 A | A H76 T | 5 | 38.5 | 1.0 | 38.7 |
| S H15 P | Q H14 K | 5 | 38.5 | 4.0 | 9.6 |
| S H15 P | S H40 G | 5 | 38.5 | 3.6 | 10.7 |
| S H15 P | Y H57 W | 5 | 38.5 | 5.3 | 7.2 |
| D H36 S | T H68 N | 5 | 100.0 | 1.9 | 52.0 |
| D H36 S | L H80 S | 5 | 38.5 | 0.8 | 47.1 |

Table S8 continued.

| | | | | Count | correlated frequ. (%) | uncorrelated frequ. (%) | enrichment |
|---|---|---|---|---|---|---|---|
| D H36 S | D H90 E | | | 5 | 100.0 | 3.8 | 26.0 |
| D H36 S | T H100 A | | | 5 | 100.0 | 5.4 | 18.6 |
| T H68 N | L H80 S | | | 5 | 100.0 | 5.4 | 18.5 |
| T H68 N | D H90 E | | | 5 | 100.0 | 25.5 | 3.9 |
| T H68 N | T H100 A | | | 5 | 100.0 | 35.7 | 2.8 |
| L H80 S | D H90 E | | | 5 | 100.0 | 10.8 | 9.2 |
| L H80 S | T H100 A | | | 5 | 100.0 | 15.2 | 6.6 |
| D H90 E | T H100 A | | | 5 | 100.0 | 71.4 | 1.4 |
| S H40 G | Y H57 W | | | 5 | 35.7 | 0.3 | 108.5 |
| I H101 V | L H21 I | T H96 S | | 14 | 93.3 | 0.42 | 224 |
| L H21 I | T H35 S | T H96 S | | 11 | 52.4 | 0.09 | 573 |
| I H101 V | T H35 S | T H96 S | | 11 | 73.3 | 0.66 | 110 |
| I H101 V | L H21 I | T H35 S | | 11 | 73.3 | 0.28 | 259 |
| L H21 I | T H96 S | V H42 I | | 5 | 21.7 | 0.05 | 474 |
| I H101 V | T H96 S | V H42 I | | 5 | 33.3 | 0.33 | 100 |
| I H101 V | L H21 I | V H42 I | | 5 | 16.1 | 0.14 | 114 |
| S H15 P | S H40 G | Y H57 W | | 5 | 38.5 | 0.25 | 153 |
| D H36 S | L H80 S | T H68 N | | 5 | 100.0 | 0.29 | 343 |
| D H36 S | D H90 E | T H68 N | | 5 | 100.0 | 1.37 | 73 |
| D H36 S | T H100 A | T H68 N | | 5 | 100.0 | 1.92 | 52 |
| D H36 S | D H90 E | L H80 S | | 5 | 100.0 | 0.58 | 171 |
| D H36 S | L H80 S | T H100 A | | 5 | 100.0 | 0.82 | 122 |
| D H36 S | D H90 E | T H100 A | | 5 | 100.0 | 3.85 | 26 |
| D H90 E | L H80 S | T H68 N | | 5 | 100.0 | 3.87 | 26 |
| L H80 S | T H100 A | T H68 N | | 5 | 100.0 | 5.41 | 18 |
| D H90 E | T H100 A | T H68 N | | 5 | 100.0 | 25.51 | 4 |
| D H90 E | L H80 S | T H100 A | | 5 | 100.0 | 10.82 | 9 |
| L H21 I | T H35 S | T H96 S | I H101 V | 11 | 73.3 | 0.042 | 1766 |
| L H21 I | V H42 I | T H96 S | I H101 V | 5 | 33.3 | 0.021 | 1599 |
| T H68 N | L H80 S | D H90 E | T H100 A | 5 | 100.0 | 3.865 | 26 |
| D H36 S | L H80 S | D H90 E | T H100 A | 5 | 100.0 | 0.583 | 171 |
| D H36 S | T H68 N | D H90 E | T H100 A | 5 | 100.0 | 1.375 | 73 |
| D H36 S | T H68 N | L H80 S | T H100 A | 5 | 100.0 | 0.292 | 343 |
| D H36 S | T H68 N | L H80 S | D H90 E | 5 | 100.0 | 0.208 | 480 |
| L H21 I | G H36 S | T H96 S | I H101 V | 4 | 26.7 | 0.036 | 742 |
| S H17 A | D H40 N | L H94 P | A H105 T | 4 | 50.0 | 0.054 | 922 |
| Q H14 K | S H15 P | S H40 G | Y H57 W | 4 | 30.8 | 0.013 | 2323 |
| T H35 S | V H42 I | T H96 S | I H101 V | 4 | 26.7 | 0.033 | 803 |
| M H39 I | D H40 N | E H64 Y | H H66 N | 4 | 44.4 | 0.139 | 319 |
| L H21 I | G H36 T | T H96 S | I H101 V | 4 | 26.7 | 0.029 | 927 |
| S H17 A | D H40 N | E H64 Y | H H66 N | 4 | 44.4 | 0.554 | 80 |
| S H17 A | M H39 I | E H64 Y | H H66 N | 4 | 36.4 | 0.147 | 247 |

Table S8 continued.

| | | | | | Count | correlated frequ. (%) | uncorrelated frequ. (%) | enrichment |
|---|---|---|---|---|---|---|---|---|
| L H21 I | T H35 S | V H42 I | I H101 V | | 4 | 26.7 | 0.014 | 1885 |
| S H17 A | M H39 I | D H40 N | H H66 N | | 4 | 44.4 | 0.627 | 71 |
| L H21 I | T H35 S | V H42 I | T H96 S | | 4 | 26.7 | 0.005 | 5837 |
| S H17 A | M H39 I | D H40 N | E H64 Y | | 4 | 44.4 | 0.331 | 134 |
| D H36 S | T H68 N | L H80 S | D H90 E | T H100 A | 5 | 100.0 | 0.208 | 480 |
| L H21 I | T H35 S | V H42 I | T H96 S | I H101 V | 4 | 26.7 | 0.002 | 12840 |
| S H17 A | M H39 I | D H40 N | E H64 Y | H H66 N | 4 | 44.4 | 0.070 | 638 |
| L H21 I | T H35 S | G H36 T | T H96 S | I H101 V | 3 | 20.0 | 0.003 | 6982 |
| L H21 I | T H35 S | G H36 S | T H96 S | I H101 V | 3 | 20.0 | 0.004 | 5586 |
| L H21 I | T H35 S | S H64 N | T H96 S | I H101 V | 3 | 20.0 | 0.007 | 2704 |
| N L56 Y | A L57 T | K L65 T | S H64 N | Q H95 H | 3 | 100.0 | 0.002 | 59701 |
| Q H14 K | S H15 P | S H40 G | Y H57 W | R H75 Q | 3 | 23.1 | 0.000 | 108457 |
| L H21 I | G H36 T | V H42 I | T H96 S | I H101 V | 3 | 20.0 | 0.001 | 13908 |
| S H17 A | M H39 I | D H40 N | E H64 Y | L H94 P | 3 | 33.3 | 0.006 | 5450 |
| S H17 A | M H39 I | D H40 N | E H64 Y | A H105 T | 3 | 37.5 | 0.041 | 913 |
| S H17 A | M H39 I | D H40 N | H H66 N | L H94 P | 3 | 33.3 | 0.012 | 2877 |
| S H17 A | M H39 I | D H40 N | H H66 N | A H105 T | 3 | 37.5 | 0.078 | 482 |
| S H17 A | M H39 I | D H40 N | L H94 P | A H105 T | 3 | 37.5 | 0.007 | 5493 |
| S H17 A | D H40 N | E H64 Y | H H66 N | L H94 P | 3 | 33.3 | 0.010 | 3258 |
| S H17 A | D H40 N | E H64 Y | H H66 N | A H105 T | 3 | 37.5 | 0.069 | 546 |
| S H17 A | E H64 Y | H H66 N | L H94 P | A H105 T | 3 | 30.0 | 0.003 | 11197 |
| M H39 I | D H40 N | E H64 Y | H H66 N | L H94 P | 3 | 33.3 | 0.003 | 12945 |
| M H39 I | D H40 N | E H64 Y | H H66 N | A H105 T | 3 | 37.5 | 0.017 | 2169 |
| M H39 I | E H64 Y | H H66 N | L H94 P | A H105 T | 3 | 30.0 | 0.001 | 44493 |
| D H40 N | E H64 Y | H H66 N | L H94 P | A H105 T | 3 | 37.5 | 0.003 | 14774 |
| Q H3 K | D H36 S | Y H57 N | T H68 N | L H80 S | 3 | 60.0 | 0.002 | 32299 |
| Q H3 K | D H36 S | Y H57 N | T H68 N | D H90 E | 3 | 60.0 | 0.009 | 6851 |
| Q H3 K | D H36 S | Y H57 N | T H68 N | T H100 A | 3 | 60.0 | 0.012 | 4894 |
| Q H3 K | D H36 S | Y H57 N | L H80 S | D H90 E | 3 | 60.0 | 0.004 | 16149 |
| Q H3 K | D H36 S | Y H57 N | L H80 S | T H100 A | 3 | 60.0 | 0.005 | 11535 |
| Q H3 K | D H36 S | Y H57 N | D H90 E | T H100 A | 3 | 60.0 | 0.025 | 2447 |
| Q H3 K | Y H57 N | T H68 N | L H80 S | D H90 E | 3 | 60.0 | 0.025 | 2437 |
| Q H3 K | Y H57 N | T H68 N | L H80 S | T H100 A | 3 | 60.0 | 0.034 | 1741 |
| Q H3 K | T H68 N | L H80 S | D H90 E | T H100 A | 3 | 60.0 | 0.302 | 198 |
| D H36 S | Y H57 N | T H68 N | L H80 S | D H90 E | 3 | 60.0 | 0.017 | 3539 |
| D H36 S | Y H57 N | T H68 N | L H80 S | T H100 A | 3 | 60.0 | 0.024 | 2528 |
| Y H57 N | T H68 N | L H80 S | D H90 E | T H100 A | 3 | 60.0 | 0.315 | 191 |

**Table S9.** Clusters of correlated convergent mutations.

| Mutations | | Shortest pairwise distances (Å) |
|---|---|---|
| T L7 S | T L8 P | 2.9 |
| E H90 Q | L H91 F | 3.7 |
| H L115 L | F L116 W | 3.7 |
| T H105 S | R H106 G | 3.7 |
| D L1 E | V L2 L | 3.7 |
| Q L3 E | M L4 L | 3.8 |
| Q H5 L | Q H6 E | 3.8 |
| V L2 I | L L3 V | 3.8 |
| Q H5 V | Q H6 E | 3.8 |
| A H105 T | R H106 L | 3.8 |
| S L92 N | S L93 T | 3.8 |
| N H66 T | Y H67 N | 3.8 |
| K H5 Q | Q H6 E | 3.8 |
| Q L3 V | M L4 L | 3.8 |
| S H64 R | T H65 I | 3.8 |
| Q H6 E | P H7 S | 3.8 |
| K H72 R | S H74 G | 3.8 |
| V H42 I | K H43 R | 3.8 |
| Q H5 E | Q H6 E | 3.8 |
| A H105 V | R H106 T | 3.8 |
| A H100 G | M H101 I | 3.8 |
| T H65 I | Y H66 H | 3.8 |
| Q H14 K | S H15 P | 3.8 |
| M H39 F | H H40 Q | 3.8 |
| L H50 F | K H51 E | 3.8 |
| Y H38 F | M H39 F | 3.8 |
| S H64 Y | T H65 I | 3.8 |
| M L4 L | S L5 T | 3.8 |
| G H63 D | S H64 N | 3.8 |
| S H35 R | S H36 N | 3.8 |
| I H2 V | Q H3 K | 3.8 |
| Y H37 F | T H38 W | 3.8 |
| A H84 V | S H85 R | 3.8 |
| L H4 V | A H25 V | 4.5 |
| T H35 S | P H58 C | 4.6 |
| W L107 R | P L116 L | 4.8 |
| N H40 H | A H105 T | 4.9 |
| S H57 N | S H64 Y | 4.9 |
| Y L38 H | G L107 S | 5.0 |
| A H17 T | S H93 R | 5.1 |
| S H57 N | S H64 N | 5.1 |
| A H54 G | Y H66 D | 5.3 |

Table S9 continued.

| Mutations | | Shortest pairwise distances (Å) |
|---|---|---|
| H L31 Y | N L34 D | 5.3 |
| G H54 A | S H66 T | 5.4 |
| T H29 S | S H36 N | 5.4 |
| S H36 G | S H58 N | 5.4 |
| M H39 I | A H105 T | 5.4 |
| M H39 I | A H105 V | 5.5 |
| S H57 T | S H64 T | 5.5 |
| F H30 L | S H36 T | 5.5 |
| A H25 T | S H85 N | 5.5 |
| S L28 T | V L30 L | 5.5 |
| T H35 I | Y H37 N | 5.6 |
| T H55 M | T H65 S | 5.6 |
| T H35 S | Y H37 F | 5.6 |
| T H29 S | T H35 S | 5.7 |
| N L34 D | N L36 D | 5.7 |
| N H40 S | R H106 Y | 5.8 |
| Y H37 F | S H57 N | 5.9 |
| N H40 H | R H106 S | 5.9 |
| Y H37 F | R H106 G | 6.0 |
| Y H37 F | P H58 T | 6.1 |
| L L30 F | Y L38 F | 6.1 |
| Y H103 F | A H105 V | 6.1 |
| E H1 K | Q H3 K | 6.1 |
| Q H3 K | K H5 Q | 6.1 |
| Q H1 E | Q H3 K | 6.1 |
| Y H37 F | R H106 T | 6.2 |
| M L4 L | S L26 G | 6.2 |
| S H63 T | T H65 I | 6.4 |
| V L2 I | M L4 L | 6.4 |
| E H1 Q | K H3 Q | 6.4 |
| V L101 I | Y L103 F | 6.4 |
| Y H103 F | T H105 S | 6.4 |
| K H90 E | N H92 D | 6.5 |
| V L3 Q | S L5 T | 6.5 |
| D H36 G | N H59 Y | 6.5 |
| I L2 L | L L4 M | 6.6 |
| V H42 L | Q H44 K | 6.7 |
| Y H37 F | R H106 S | 6.7 |
| S H15 P | S H93 N | 6.8 |
| Q H6 E | L H21 I | 7.0 |
| S H66 T | N H68 S | 7.1 |
| F H35 S | D H59 N | 7.4 |
| V H42 I | A H105 T | 7.4 |

Table S9 continued.

| Mutations | | Shortest pairwise distances (Å) |
|---|---|---|
| T H35 S | S H62 T | 7.5 |
| V H80 A | S H85 N | 7.6 |
| K H70 R | G H74 D | 7.6 |
| T H65 I | I H78 V | 7.6 |
| I H56 F | V H80 A | 7.6 |
| S L108 T | P L116 L | 7.7 |
| A H58 S | S H64 N | 7.8 |
| T H35 S | S H85 N | 7.8 |
| M L94 L | A L99 G | 8.0 |
| F H30 L | L H87 V | 8.1 |
| S H57 R | S H66 R | 8.1 |
| T H25 A | A H84 T | 8.2 |
| V L30 L | N L34 D | 8.2 |
| H H40 Y | Y H103 F | 8.2 |
| T H25 A | T H29 S | 8.4 |
| T H29 I | S H85 R | 8.4 |
| G H63 D | K H66 E | 8.5 |
| S L22 T | T L90 S | 8.6 |
| T H29 S | S H85 N | 8.6 |
| S H36 T | K H82 R | 8.6 |
| S H35 N | S H57 N | 8.7 |
| Y H57 N | E H66 K | 8.7 |
| S H36 R | S H57 T | 8.8 |
| G H63 D | L H78 M | 8.8 |
| Y L38 F | S L109 T | 8.8 |
| Y L38 F | E L109 D | 8.8 |
| T L10 S | V L101 I | 8.8 |
| N L40 S | R L66 K | 8.8 |
| K H66 E | L H78 M | 8.8 |
| S L92 N | Q L95 K | 8.9 |
| G H27 E | S H85 N | 9.0 |
| Y H37 F | I H56 V | 9.0 |
| S H36 T | N H82 I | 9.0 |
| D H55 N | A H105 T | 9.0 |
| M H39 I | V H80 A | 9.1 |
| A H9 P | M H21 I | 9.1 |
| S H66 T | K H72 R | 9.2 |
| A H25 T | R H106 T | 9.2 |
| N H57 D | S H66 N | 9.3 |
| M L4 L | T L7 S | 9.3 |
| Y H37 N | A H105 G | 9.3 |
| T H55 A | A H105 T | 9.3 |
| T H35 I | M H39 I | 9.3 |

Table S9 continued.

| Mutations | | | Shortest pairwise distances (Å) | | |
|---|---|---|---|---|---|
| T H25 A | T H105 S | | 9.3 | | |
| L L9 S | V L101 I | | 9.4 | | |
| Y H55 F | A H105 T | | 9.4 | | |
| M H39 I | Y H88 F | | 9.4 | | |
| M H39 I | T H65 S | | 9.5 | | |
| Y H37 N | A H105 V | | 9.5 | | |
| Y H3 Q | Q H6 E | | 9.6 | | |
| Q H3 K | Q H6 E | | 9.6 | | |
| A H25 V | A H105 T | | 9.7 | | |
| T H35 S | M H39 I | | 9.7 | | |
| K H3 Q | E H6 Q | | 9.7 | | |
| A H25 T | R H106 G | | 9.7 | | |
| K H3 Q | T H105 S | | 9.7 | | |
| K H5 Q | A H105 V | | 9.8 | | |
| L L4 M | S L7 T | | 9.8 | | |
| Q H3 K | A H105 T | | 9.8 | | |
| S L28 T | Q L106 H | | 9.9 | | |
| I H2 V | V H5 Q | | 9.9 | | |
| Y H62 S | S H66 N | | 9.9 | | |
| S L37 G | P L116 L | | 9.9 | | |
| E H51 D | K H70 R | | 9.9 | | |
| D L1 E | V L2 L | L L3 V | 3.8 | 3.8 | 5.9 |
| Q L1 E | I L2 L | L L4 M | 3.8 | 6.6 | 9.6 |
| Y H38 W | N H40 H | A H105 T | 5.3 | 6.5 | 8.5 |
| Q H5 E | Q H6 E | Y H103 F | 3.8 | 8.0 | 10.0 |
| D L1 E | V L2 L | S L107 G | 3.7 | 10.2 | 10.3 |
| A H25 T | G H27 E | D H36 E | 5.8 | 9.7 | 10.5 |
| Q H3 K | Q H6 E | A H105 V | 9.8 | 10.2 | 10.5 |
| Q H3 K | Q H6 E | S H85 N | 10.4 | 11.3 | 11.7 |
| E H1 Q | K H3 Q | V H5 Q | 6.7 | 6.7 | 13.4 |
| Q H3 K | Q H6 E | P H7 S | 3.8 | 9.8 | 12.8 |
| Q H6 E | P H7 S | M H89 I | 3.8 | 10.9 | 12.3 |
| A H25 T | N H40 H | A H105 T | 5.3 | 9.7 | 12.7 |
| T H35 I | M H39 I | A H105 T | 6.0 | 9.3 | 13.9 |
| M H39 I | M H89 I | A H105 T | 5.8 | 10.2 | 12.1 |
| M H39 I | S H85 N | A H105 T | 5.8 | 10.7 | 13.5 |
| I L2 L | L L4 M | S L7 T | 6.6 | 9.9 | 16.4 |
| T H29 S | T H35 S | M H39 I | 6.2 | 10.4 | 13.7 |
| A H25 T | M H39 I | I H56 L | 5.1 | 11.8 | 15.5 |
| T H25 A | V H42 I | T H105 S | 8.0 | 9.3 | 16.1 |
| T H29 S | T H35 S | S H64 N | 5.9 | 11.7 | 17.1 |
| T H35 S | S H64 N | N H66 H | 7.1 | 11.7 | 16.4 |
| Q H6 E | M H39 I | V H42 I | 10.2 | 11.7 | 12.3 |

Table S9 continued.

| Mutations | | | | | Shortest pairwise distances (Å) | | | | | |
|---|---|---|---|---|---|---|---|---|---|---|
| K H5 V | T H65 I | V H87 I | | | 10.3 | 11.9 | 20.4 | | | |
| E H1 Q | Q H6 E | A H105 V | | | 10.6 | 11.7 | 16.0 | | | |
| A H25 T | D H36 E | Y H38 T | N H40 H | | 6.6 | 6.6 | 9.7 | 11.4 | 12.5 | 12.6 |
| E H1 Q | K H3 Q | E H5 Q | R H106 G | | 6.2 | 6.4 | 10.5 | 11.7 | 12.4 | 14.1 |
| A H25 T | Y H38 T | N H40 H | S H85 T | | 5.5 | 6.6 | 11.4 | 11.4 | 12.6 | 13.9 |
| Q H6 E | M H39 I | S H85 N | M H89 I | | 10.2 | 10.7 | 11.7 | 12.0 | 12.7 | 13.9 |
| S H17 A | M H39 I | D H40 N | E H64 Y | H H66 N | 3.7 | 7.0 | 9.3 | 10.4 | 10.8 | 12.0 |
| Q H14 K | S H15 P | S H40 G | Y H57 W | R H66 Y | 3.8 | 8.8 | 8.8 | 9.3 | 24.1 | 24.5 |
| A H9 P | M H39 I | D H64 N | A H65 T | V H101 I | 3.8 | 9.0 | 9.6 | 10.6 | 16.4 | 19.1 |
| A H9 P | M H39 I | A H65 T | S H66 N | V H101 I | 3.7 | 9.0 | 9.6 | 10.8 | 16.4 | 19.1 |
| A H9 P | M H39 I | A H65 T | M H89 L | V H101 I | 9.0 | 9.5 | 10.5 | 11.5 | 12.2 | 13.8 |
